# Intercellular Conduction Optimizes Arterial Network Function and Conserves Blood Flow Homeostasis during Cerebrovascular Challenges

**DOI:** 10.1101/625525

**Authors:** Anil Zechariah, Cam Ha T. Tran, Bjorn O. Hald, Shaun L. Sandow, Maria Sancho, Sergio Fabris, Ursula I. Tuor, Grant R.J. Gordon, Donald G. Welsh

## Abstract

Cerebral arterial networks match blood flow delivery with neural activity. Neurovascular response begins with a stimulus and a focal change in vessel diameter, which by themselves is inconsequential to blood flow magnitude, until they spread and alter the contractile status of neighboring arterial segments. We sought to define the mechanisms underlying integrated vascular behavior and considered the role of intercellular electrical signalling in this phenomenon. Electron microscopic and histochemical analysis revealed the structural coupling of cerebrovascular cells and the expression of gap junctional subunits at the cell interfaces, enabling intercellular signaling among vascular cells. Indeed, robust vasomotor conduction was detected in human and mice cerebral arteries after focal vessel stimulation; a response attributed to endothelial gap junctional communication, as its genetic alteration attenuated this behavior. Conducted responses was observed to ascend from the penetrating arterioles, influencing the contractile status of cortical surface vessels, in a simulated model of cerebral arterial network. Ascending responses recognised *in vivo* after whisker stimulation, were significantly attenuated in mice with altered endothelial gap junctional signalling confirming that gap junctional communication drives integrated vessel responses. The diminishment in vascular communication also impaired the critical ability of the cerebral vasculature to maintain blood flow homeostasis and hence tissue viability, after stroke. Our findings establish the integral role of intercellular electrical signalling in transcribing focal stimuli into coordinated changes in cerebrovascular contractile activity and expose, a hitherto unknown mechanism for flow regulation after stroke.

**Significance:** Neurovascular responses are viewed as a one step process whereby stimuli derived from neural cells focally diffuse to a neighboring vessel, altering its contractile state. While focal changes in tone can subtly tune flow distribution, they can’t substantively change “perfusion magnitude” as vascular resistance is broadly distributed along the cerebral arterial tree. We report that nature overcomes this biophysical constraint by conducting electrical signals among coupled vascular cells, along vessels, and across branch points. Our quantitative exploration of intercellular conduction illustrates how network coordination optimizes blood flow delivery in support of brain function. Diminishing the ability of vascular cells to electrically communicate, mitigates the brain’s ability to regulate perfusion during functional hyperemia and after stroke, the latter advancing tissue injury.

## Introduction

Cerebral arteries are linked in series and parallel, forming an integrated network responsible for matching blood flow delivery with energetic demand (1). The coupling process is commonly perceived as beginning with neural cells generating a stimulus and ending with its diffusion to a neighboring blood vessel, altering the contractile state of vascular smooth muscle (2). While focal changes in vasomotor tone can subtly tune blood flow distribution, they are unable to initiate larger changes in perfusion magnitude, which is needed to support robust neural activity. The latter response will only occur if vascular resistance changes in multiple arterial segments, upstream from the focal site of neural stimulation.

As vascular resistance is broadly distributed in network structures, the arterial wall must be encoded with mechanism(s) to convert focal neural stimuli into broader multi-segmental responses. The identity of these mechanisms is uncertain and a center point of investigation in the vascular field. Of note in the peripheral vasculature is the ability of varied but focal stimuli, to elicit responses that spread along arteries and across branch points (3, 4). The so called “conducted response” is independent of metabolite diffusion, hemodynamics and innervation, and requires a common signal to spread among thousands of vascular cells (5-7). The movement of ions (charge) via vascular gap junctions is that signal and it coordinates changes in membrane potential (V_M_), cytosolic [Ca^2+^] and myosin light chain phosphorylation (8), altering vessel contractility. Vascular gap junctions are comprised of two hemi-channels of six connexin (Cx) subunits (i.e. Cx37 (9-12), Cx40 (10-13) Cx43 (12,14) and Cx45 (14-17); each expressed at highly variable levels, depending on the type of cells they connect (18,19).

The process of intercellular conduction has escaped deep quantitative examination in the brain, a knowledge gap that compromises understanding of integrated blood flow control, in this critical organ. Consequently, we crafted a clear quantitative framework to rationalize conduction, its electrical underpinnings and how they integrate into and drive the control of brain blood flow. Foundational experiments focused on arterial structure, examining intercellular connections and Cx40 abundance, a key gap junctional subunit exclusive to endothelial cells, in mice (8, 20-22). In the *ex vivo* setting, robust conduction was observed, and this response was impaired upon the elimination of endothelium or Cx40. A virtual network, built from structural and electrical observations, revealed that the arterial wall is optimally designed to facilitate the “ascension” of vascular responses from penetrating arterioles into the cortical surface network. Multi-photon imaging of awake mice detected ascension after whisker stimulation, and deletion of Cx40 strikingly impaired the upward spread of vessel dilation. The process of ascension need not be limited to functional hyperemic responses but could, as in the case of stroke, aid in blood flow homeostasis and neuroprotection. This relationship was confirmed *in vivo*, with the deletion of Cx40 blunting the compensatory blood flow responses to stroke, expanding tissue injury. Vascular electrical signalling is integral in translating discrete neural events into broader multi-segmental responses, necessary to support neurovascular coupling and to maintain critical levels of perfusion during severe cerebrovascular challenges.

## Results

### The cerebral arterial wall is structurally encoded for conduction

The spread of electrical signals along cerebral arteries requires intercellular contacts and the presence of gap junctions. Mouse middle cerebral arteries analyzed using electron microscopy revealed robust cellular contact sites, principally among neighboring endothelial cells (Fig. 1a,b). Contact sites were sparser among smooth muscle cells and between the two cell layers, with the latter requiring projections to form and extend through a connective tissue layer (19) (Fig. 1c,d). Endothelial-to-smooth muscle projections are particularly striking given the thickness of the internal elastic lamina (IEL) and the distance between the two cell layers. Contact sites irrespective of the location were electron dense with regions of pentalaminar membrane, suggestive of Cx expression and the presence of gap junctions (19). In this regard, Cx40 was detected between longitudinally oriented endothelial cells (Fig. 1e,f; arrowheads, g), indicative of its potential role in longitudinal charge spread. Note, punctate Cx40 was observed near ‘IEL holes’ (Fig. 1f (arrow)), consistent with Cx40 localization at the: 1) endothelial-smooth muscle interface; or 2) in the endoplasmic reticulum, found at the base of projections. Cx40 labelling was absent; in control tissues treated with secondary antibody alone or with a Cx40 peptide that competes for the primary antibody binding site (Fig. 1h(inset)) and in the smooth muscle layer (Fig.1i, j). Subsequently, Cx40 expression was also confirmed in the endothelial layer of human cerebral arteries (“SI Appendix, Fig.1”). A broader immunohistochemical investigation revealed the abundance of Cx37, Cx40 and Cx43 in the endothelial cell layer of mouse middle cerebral arteries (“SI Appendix, Fig.2a-c”). Punctate Cx37 and Cx43 labelling was also observed near IEL holes indicating its localization near the endothelial-smooth muscle interface (“SI Appendix, Fig.2a, c”).

**Figure 1:**
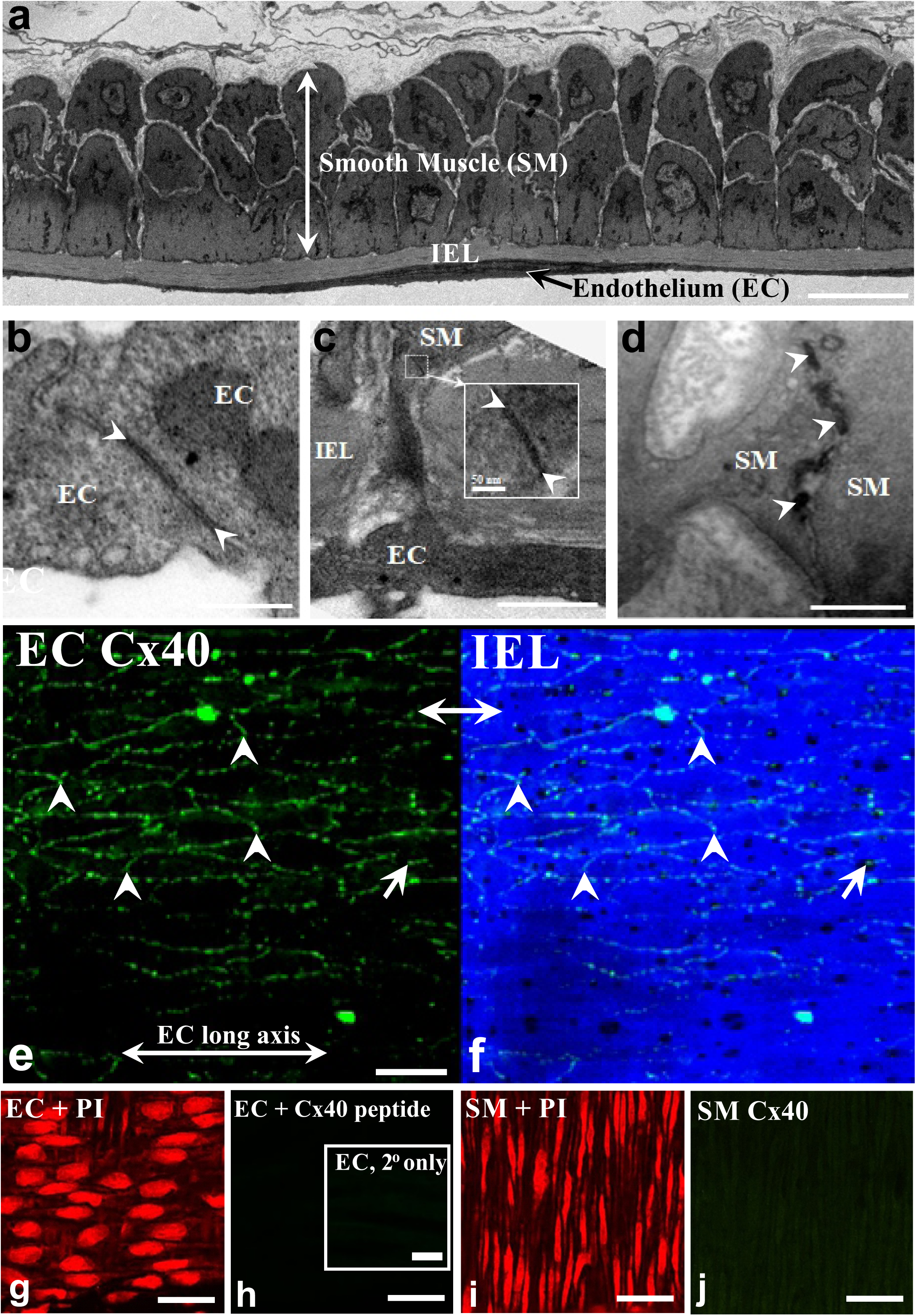
Vascular cells in cerebral arteries connect to one another and express Cx40. **(a)** Electron microscopy reveals that mouse middle cerebral arteries are composed of smooth muscle (SM), internal elastic lamina (IEL) and endothelial cell (EC) layers. Bar = 5 µm. **(b-d)** Contact sites (arrowheads) between endothelial cells (**b**, bar = 0.25 µm), between smooth muscle and endothelium (**c**, scale bar 0.25 µm), and between smooth muscle cells (**d**, bar = 0.5 µm). **(e)** Cx40 (green), a key gap junctional protein was identified by immunohistochemistry in middle cerebral arteries isolated from mice. **(f)** IEL was delineated by 488 nm autofluorescence (**g**,**i)** Propidium iodide labeling verified EC and SMC patency. (**h**) Endothelial Cx40 signal was absent in sections incubated without the primary antibody (inset) and in endothelial cells treated with a Cx40 peptide that competes for the primary antibody binding site. (**j**) Cx40 staining was also absent in the smooth muscle cell layer. Bar, e-j = 25 µm

### Endothelial/gap junctional communication drives conducted responses

Mouse and human cerebral arteries (∼100 µm diameter) were mounted in a pressure myograph for *ex vivo* conduction experiments (Fig. 2a). The discrete application of a depolarizing stimulus (high pipette [K^+^]) to one end of a human or a mouse (C57BL/6) cerebral artery, elicited a focal constriction that robustly conducted (within ∼300ms) to the distal 1800 μm site in an electrotonic manner (Fig. 2b, c). Consistent with the endothelium operating as the primary pathway for longitudinal charge spread (7), air bubble disruption eliminated conduction beyond the focal site of stimulation (Fig. 2d). Genetic ablation of Cx40 diminished but did not eliminate conducted vasomotor responses (Fig. 2c; middle column) consistent with an increase in endothelial-to-endothelial coupling resistance. We subsequently calculated the change in endothelial-to-endothelial coupling resistance using a validated model of electrical communication that accounted for the structural, gap junctional and ionic properties of vascular cells. The latter entailed incorporating polynomial fits of IV curves from isolated endothelial and smooth muscle cells (“SI Appendix, Fig. 3”). A nonlinear voltage-diameter relationship generated from cerebral arterial observations was used to convert V_M_ into a vasomotor response (23) (“SI Appendix, Fig. 4”). This approach yielded endothelial-to-endothelial coupling resistances of 1.7 MΩ and 6 MΩ in C57BL/6 and Cx40^−/−^ mice, respectively (Fig. 2c, right column). Our modelling data also predicted the absence of conduction in endothelial denuded arteries (Fig. 2d, right column) and that the rise in endothelial-to-endothelial coupling resistance in Cx40^−/−^ arteries would starkly diminish electrical spread and vasomotor response, a 1000 μm from the point of stimulation (“SI Appendix, Fig. 5”). Arterial V_M_ measurements confirmed that spreading electrical responses were indeed attenuated, 900 μm from the site of agent application in Cx40^−/−^ arteries (Fig 2e). A subset of experiments also revealed that hyperpolarizing stimuli (lower pipet [K^+^], acetylcholine) could induce a small but consistent conducted vasodilation that decayed along surface vessels and penetrating arterioles (Fig. 3a-c). Cx40 expression was also observed in the endothelial cell layer of penetrating arterioles in mice (Fig. 3d).

**Figure 2:**
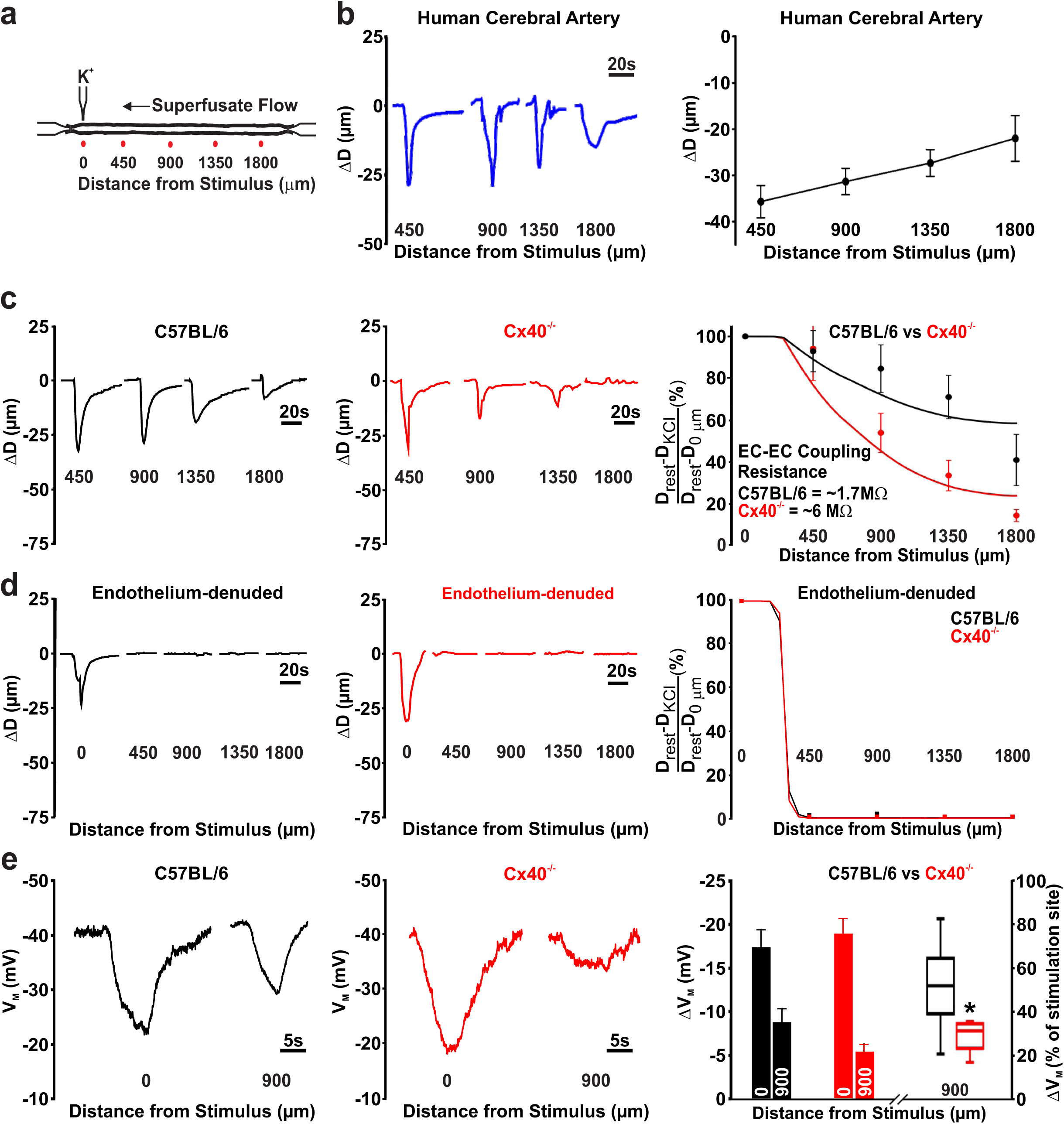
Conducted responses in cerebral arteries require intact endothelium and gap junctions. **(a)** General conduction paradigm: An isolated, pressurised middle cerebral artery was stimulated by pressure injecting KCl (500 mM, 10s, 30 PSI) through a micropipette; vasomotor responses monitored 0, 450, 900, 1350 and 1800 µm, distal to the point of stimulation. **(b)** K^+^ induced conducted responses were observed in cerebral arteries isolated from resected human tissue. **(c)** Genetic deletion of Cx40 attenuated K^+^ induced conduction in mouse cerebral arteries. Conducted responses were similar among C57BL/6 and Cx40^−/−^ mice near the point of K^+^ application (450 µm site); enhanced conduction decay was observed in Cx40^−/−^ mice at the 900 µm (U = 18.50, P<0.01, Mann-Whitney *U* test; control, n=10 vessels; Cx40^−/−^, n=10 vessels), 1350 µm (*U* = 11, P<0.01, Mann-Whitney *U* test; C57BL/6, n = 9 vessels; Cx40^−/−^, n = 10 vessels), and 1800 µm (*U* = 13, P<0.01, Mann-Whitney *U* test; C57BL/6, n = 8 vessels; Cx40^−/−^, n = 10 vessels) sites. Data in (c, right column) were fitted using a validated computational model to calculate endothelial-to-endothelial (EC-EC) coupling resistance. **(d)** The endothelium’s role in conduction was tested by passing air bubbles through the lumen to denude this layer. Data in (**d**, right column) were fitted using a validated computational model in which the endothelium had been electrically uncoupled. **(e)** The spreading membrane potential (V_M_) response in Cx40^−/−^ arteries was diminished at the conducted site (900 µm). Summary data (right column) is presented in absolute terms or with the 900 µm site expressed relative to the stimulation site (U = 4, P<0.05, Mann-Whitney *U* test; control, n=6 vessels; Cx40^−/−^, n=5 vessels).

**Figure 3.**
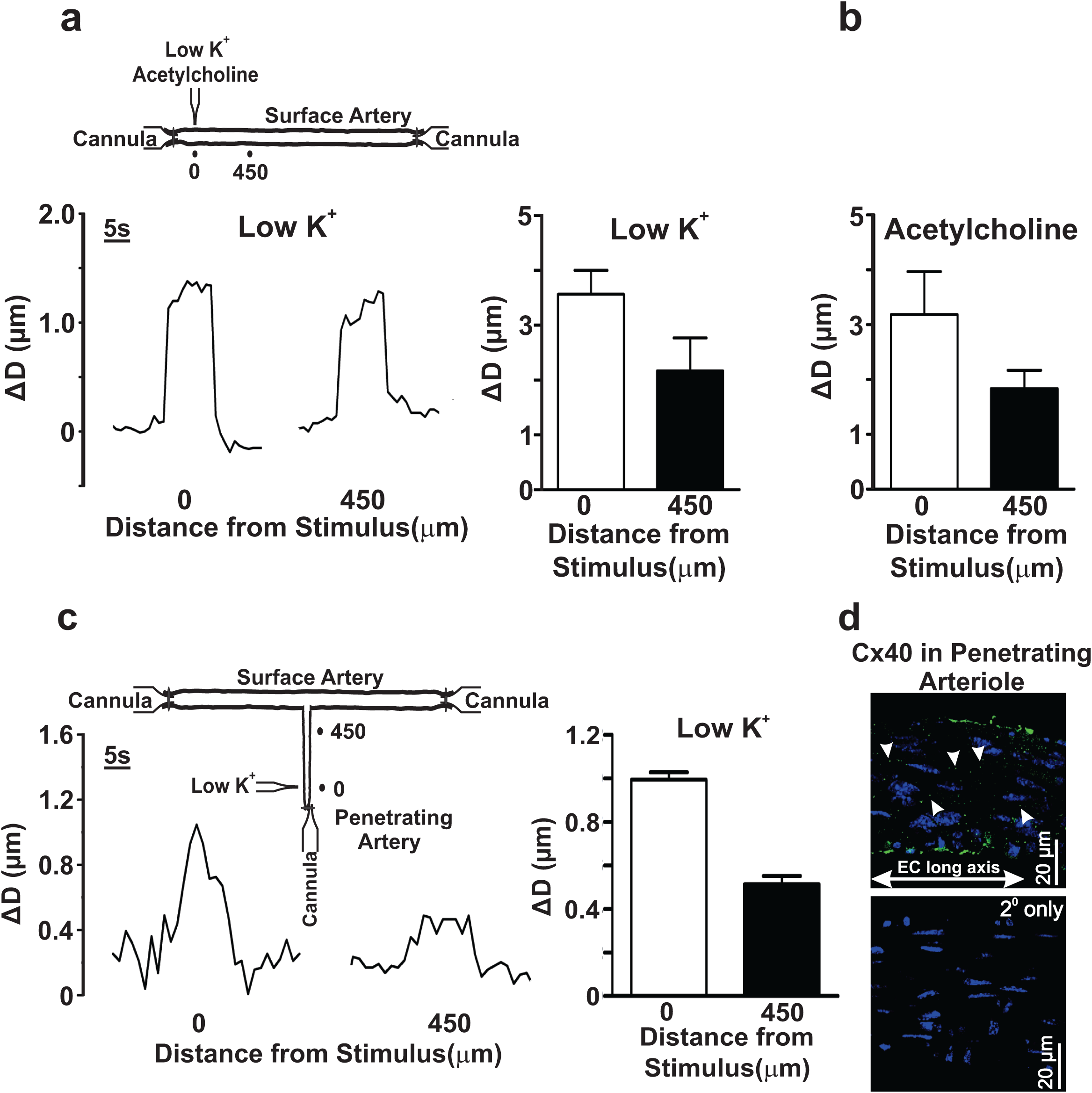
Conducted vasodilation in middle cerebral arteries and penetrating arterioles from C57BL/6 mice. **(a-b)** Middle cerebral arteries were isolated and pressurized to 50 mmHg (**Inset**). A micropipette containing ‘low pipette [K^+^]’ (25mM) or acetylcholine (1 mM) was positioned next to the arterial wall. The solution was pressure ejected (10 secs, 30 psi) and diameter was measured at the local application site (0 μm) and one conducted site (450 μm). Representative tracing and summary data of the local and conducted vasodilatory response (resting diameter for the ‘low pipette [K^+^]’ (n=4 vessels) and acetylcholine groups (n=3 vessels) were 91+14 and 76+8, respectively). **(c)** A middle cerebral artery with penetrating arteriole was isolated and triple cannulated (**Inset**). The preparation was pressurized to 50 mmHg; a pipet containing a 25 mM K^+^ saline solution was placed next to the arterial wall and pressure ejected (10 secs, 30 psi). Diameter was measured at the local (0 μm) and conducted site (450 μm). Summary data (n=4 vessels; resting diameter 47+5) of the local and conducted vasodilatory responses. **(d)** Cx40 (green) was identified by immunohistochemistry in the endothelial layer of penetrating arterioles from C57BL/6 mice.

**Figure 4:**
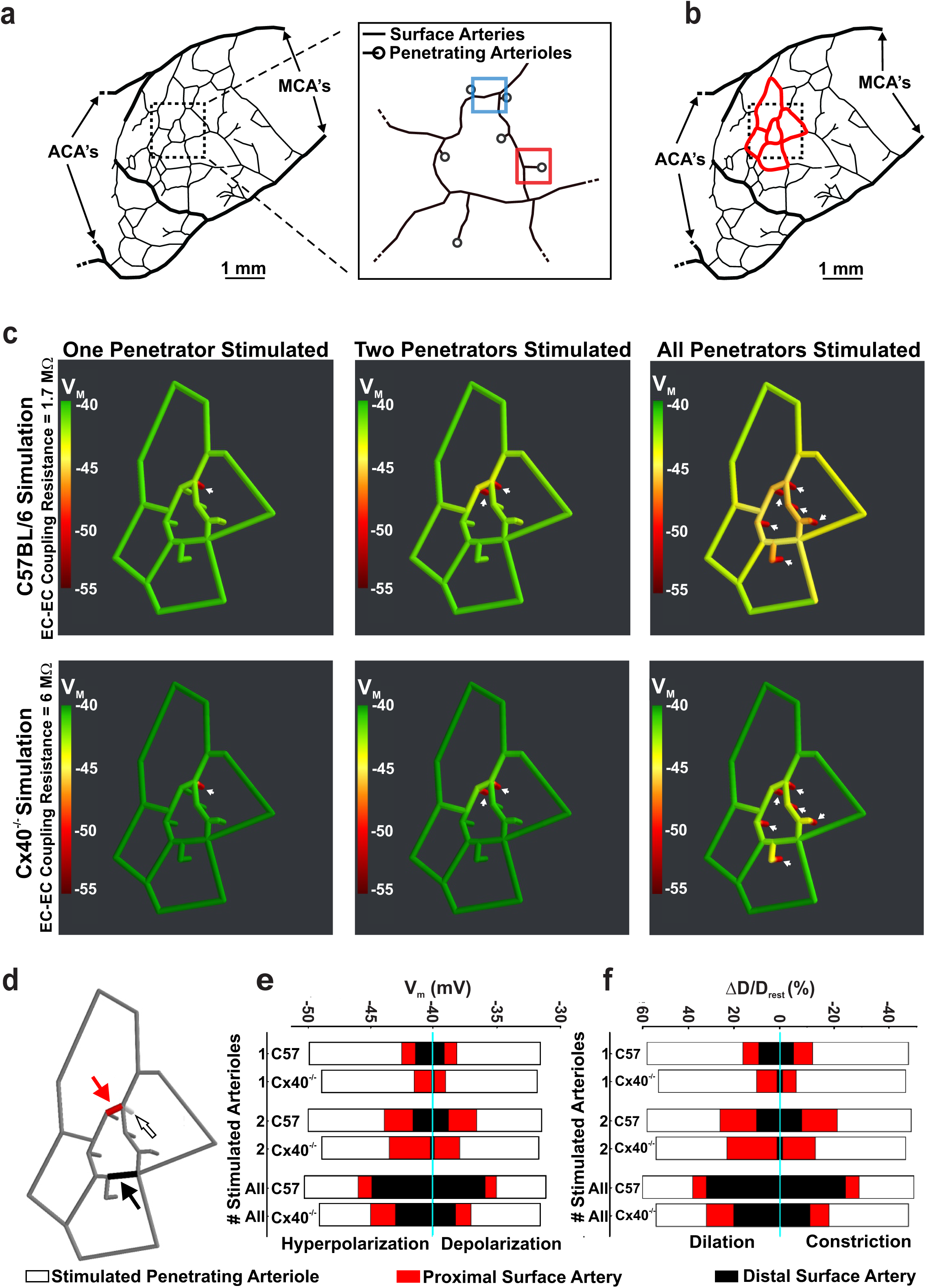
Vascular cells retain biophysical properties that support the ascension of responses from penetrating arterioles to surface vessels. **(a, b)** Using observations from Shih et al., 2009, a network model of surface arteries and penetrating arterioles was built to study the spread of electrical/vasomotor responses. The *in silico* model consisted of an interconnected network of surface arteries and six adjoining penetrating arterioles. Simulations entailed voltage clamping (15 mV positive or negative to resting V_M_, 500 ms) one distal segment in one, two or all penetrating arterioles and resolving the voltage/vasomotor response throughout the network. **(c)** The ascension of hyperpolarization was color mapped along the network; endothelial-to-endothelial coupling resistance was set to 1.7 (C57BL/6 control) or 6 (Cx40^−/−^) MΩ (calculated in Figure 2). **(d-f)** One distal segment in one, two or all penetrating arterioles was/were stimulated and steady state responses (smooth muscle V_M_ and diameter) plotted at 3 sites in the network. Those included the penetrating arteriole (200 μm upstream from the stimulation site) and two surface vessel sites. (**f**) D and D_rest_ represent peak response and resting diameters, respectively. MCA’s = Middle cerebral arteries; ACA’s = Anterior cerebral arteries. Note, copyright request for permission to publish (Fig. 3a) is currently pending.

**Figure 5:**
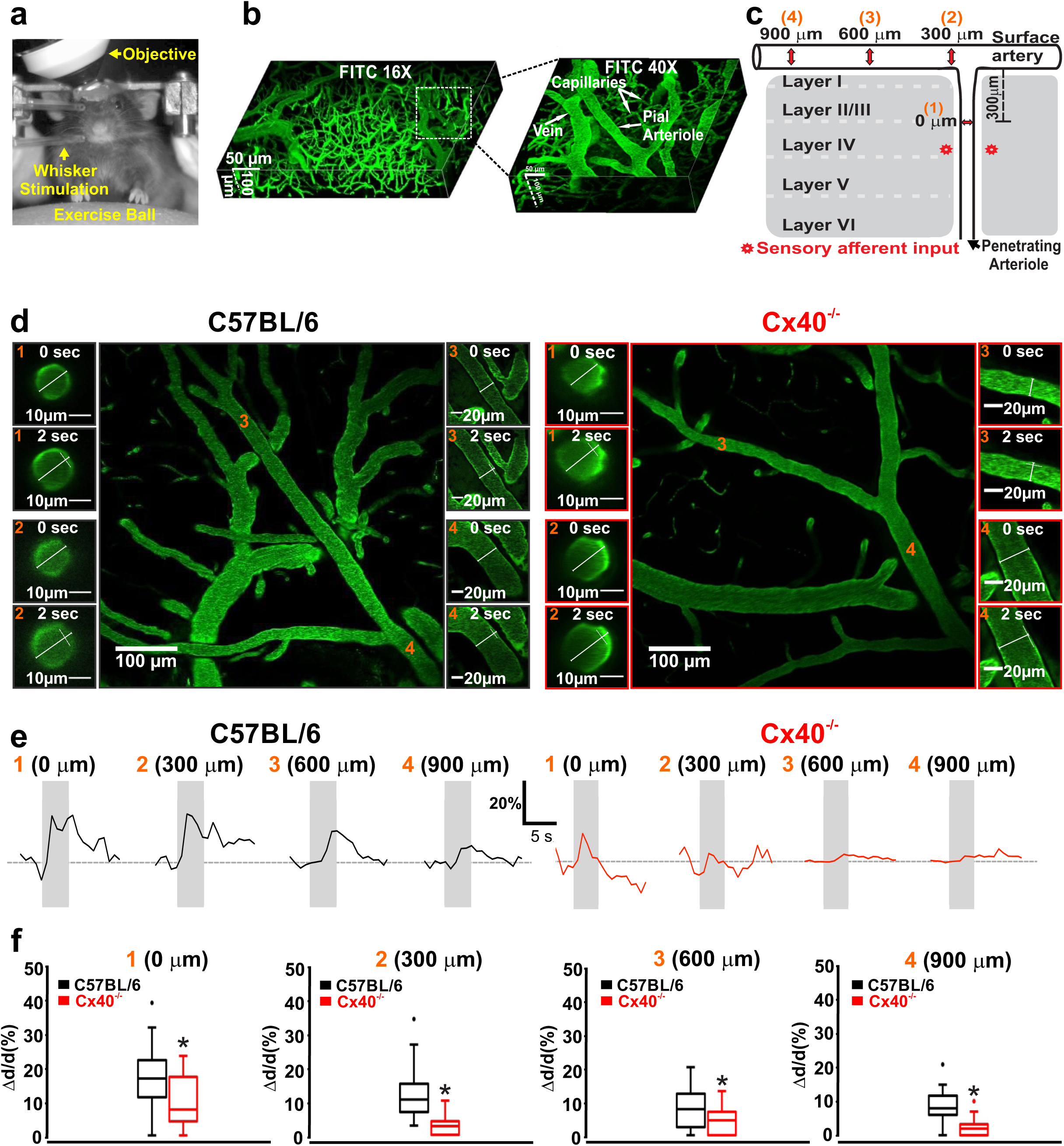
Compromised vascular conduction impairs neurovascular coupling responses. (**a**,**b**) Awake mice were positioned on an exercise ball with their head fixed to an external apparatus, to facilitate two-photon imaging of the microvasculature labeled with FITC Dextran. **c**) Vasomotor responses to whisker stimulation were monitored at 4 sites upstream (orange numbers) near to the point of barrel cortex activation. (**d**) Photomicrographs of functional dilation at four vessel sites, prior to (0 s) and during (2 s) whisker stimulation in C57BL/6 and Cx40^−/−^ mice. (**e**,**f**) Representative tracings and summary data of the functional vasomotor response to whisker stimulation. Measurements show significant attenuation of conducted responses in the penetrating arterioles and cortical surface vessels of Cx40^−/−^ mice (0 µm: (*U* = 85, P<0.05, Mann-Whitney *U* test; C57BL/6, n = 21 vessels; Cx40^−/−^, n = 14 vessels)), (300 µm: (*U* = 24, P<0.05, Mann-Whitney *U* test; C57BL/6, n = 21 vessels; Cx40^−/−^, n = 14 vessels)), (600 µm: (*U* = 126, P<0.05, Mann-Whitney *U* test; C57BL/6, n = 19 vessels; Cx40^−/−^, n = 21 vessels)), (900 µm: (*U* = 22.50, P<0.05, Mann-Whitney *U* test; C57BL/6, n = 9 vessels; Cx40^−/−^, n = 16 vessels)). * denotes significant difference between groups.

Control qPCR experiments confirmed the presence of Cx37, Cx40, Cx43 and Cx45 mRNA in human and mice cerebral arteries (“SI Appendix, Fig. 6”). Cx40 mRNA was particularly abundant in human cerebral arteries (“SI Appendix, Fig. 6a”); Immunohistochemical analysis of Cx40^−/−^ mice revealed an absence of Cx40 labeling in the endothelium and a marked reduction in Cx37 and Cx43 expression (“SI Appendix, Fig. 7a-c”). The decrease in Cx37 and Cx43 expression was surprising as quantitative PCR analysis revealed a significantly higher Cx37 mRNA and a comparable expression of Cx43 in Cx40^−/−^ mice, compared to C57BL/6 animals (“SI Appendix, Fig. 6b”). Myogenic tone and vessel reactivity to 60 mM K^+^ was similar among vessels isolated from C57BL/6 and Cx40^−/−^ mice (“SI Appendix, Fig. 8 a, b”).

**Figure 6:**
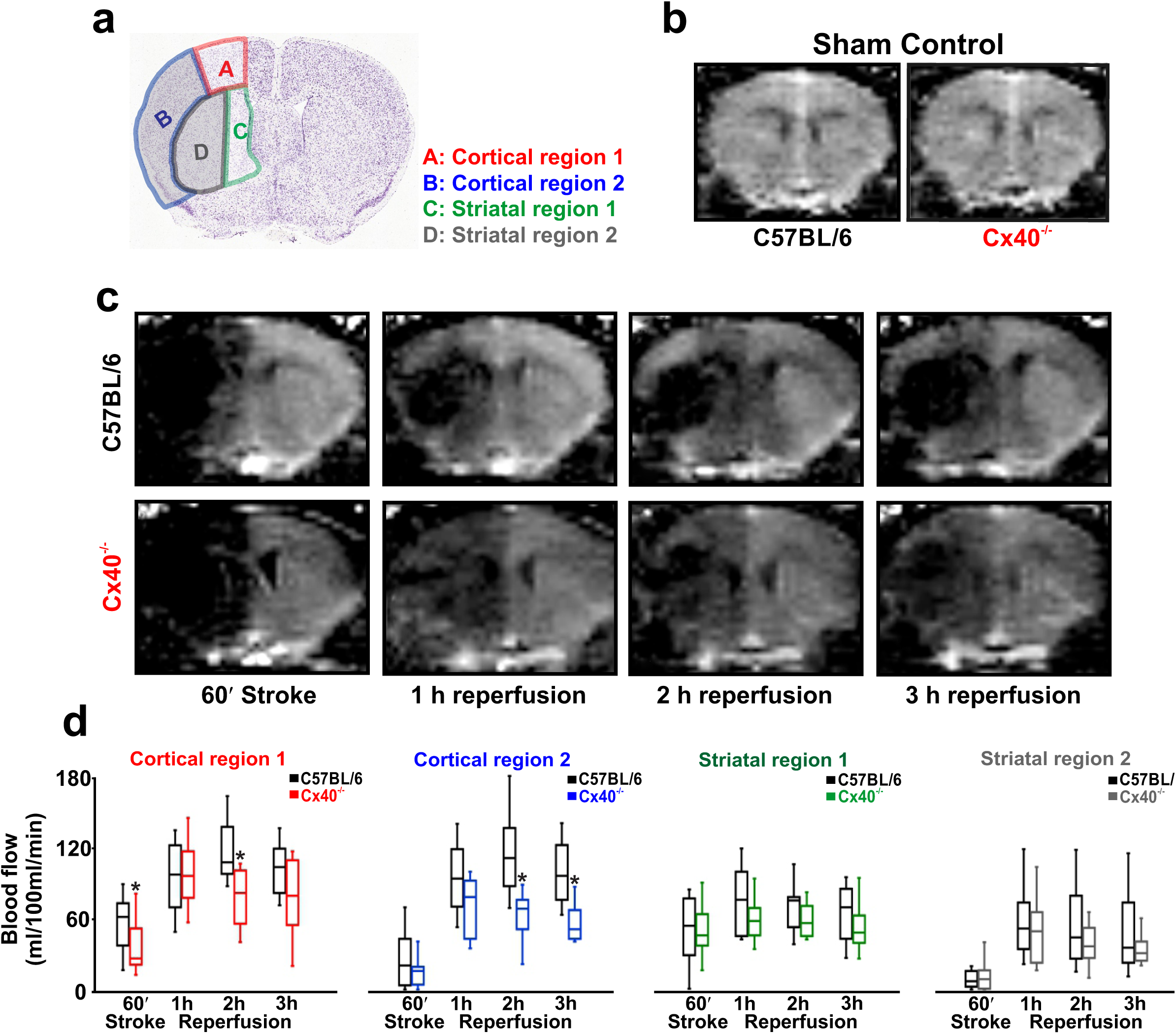
The initial blood flow response to stroke is attenuated in Cx40^−/−^ mice. **(a)** Perfusion was monitored during and following stroke in four regions of interest: cortical region 1 (A), cortical region 2 (B), striatal region 1 (C) and striatal region 2 (D) (Image credit: Allen Institute). Anatomically, regions B and D are supplied by the middle cerebral artery whereas regions A and C are watershed regions supplied by the middle, and the anterior or the posterior cerebral arteries. (**b**) ASL-MR perfusion images demonstrate that blood flow is similar in the cortex of sham controlled C57BL/6 and Cx40^−/−^ mice (*U* = 2, P>0.05, Mann-Whitney *U* test; C57BL/6, n = 4 mice; Cx40^−/−^, n = 4 mice) and in striatum (*U* = 3, P>0.05, Mann-Whitney *U* test; C57BL/6, n = 4 mice; Cx40^−/−^, n = 4 mice). (**c**) ASL-MR perfusion images from C57BL/6 and Cx40^−/−^ mice during stroke and reperfusion. (**d**) Summary data and quantitative non-parametric analysis of blood flow responses to stroke and reperfusion in C57BL/6 and Cx40^−/−^ mice. In comparison to control mice, a significant decline in blood flow was observed in Cx40^−/−^ mice, during stroke in cortical region 1: (*U* = 24, P<0.05, Mann-Whitney *U* test; C57BL/6, n = 9 mice; Cx40^−/−^, n = 10 mice), during 2 h reperfusion in cortical region 1: (*U* = 13, P<0.05, Mann-Whitney *U* test; C57BL/6, n = 9 mice; Cx40^−/−^, n = 10 mice); cortical region 2 (*U* = 5, P<0.05, Mann-Whitney *U* test; C57BL/6, n = 9 mice; Cx40^−/−^, n = 10 mice) and during 3h reperfusion in cortical region 2 (*U* = 5, P<0.05, Mann-Whitney *U* test; C57BL/6, n = 9 mice; Cx40^−/−^, n = 10 mice). * denotes significant difference between groups.

**Figure 7:**
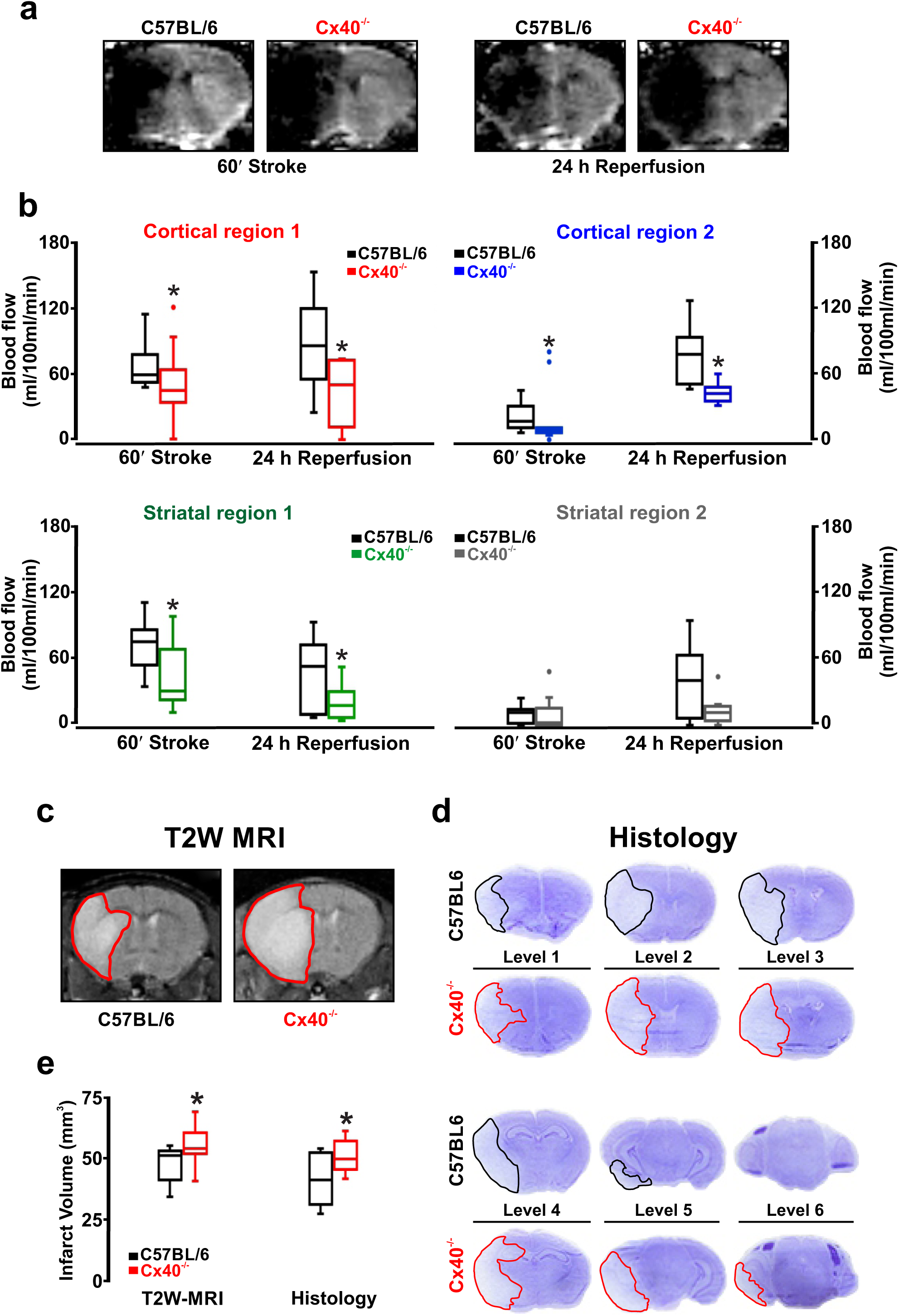
The prolonged blood flow response to stroke is attenuated in Cx40^−/−^ mice, augmenting tissue injury. Perfusion was monitored by ASL-MR during stroke (60 min) and reperfusion (24 h) in four regions of interest: cortical region 1&2 and striatal region 1&2 (see Figure 5). (**a**) ASL-MR perfusion images from C57BL/6 and Cx40^−/−^ mice during stroke and 24 h reperfusion. (**b**) Summary data and quantitative non-parametric analysis of blood flow responses to stroke and 24h reperfusion in C57BL/6 and Cx40^−/−^ mice. A significant decline in blood flow was observed in Cx40^−/−^ mice, during stroke in cortical region 1 (*U* = 21, P<0.05, Mann-Whitney *U* test; C57BL/6, n = 9 mice; Cx40^−/−^, n = 9 mice); in cortical region 2 (*U* = 17, P<0.05, Mann-Whitney *U* test; C57BL/6, n = 9 mice; Cx40^−/−^, n = 9 mice); in striatal region 1 (*U* = 21, P<0.05, Mann-Whitney *U* test; C57BL/6, n = 9 mice; Cx40^−/−^, n = 9 mice) and during 24 h reperfusion in cortical region 1 (*U* = 17, P<0.05, Mann-Whitney *U* test; C57BL/6, n = 9 mice; Cx40^−/−^, n = 9 mice); in cortical region 2 (*U* = 6, P<0.05, Mann-Whitney *U* test; C57BL/6, n = 9 mice; Cx40^−/−^, n = 9 mice) and in striatal region 1 (*U* = 20, P<0.05, Mann-Whitney *U* test; C57BL/6, n = 9 mice; Cx40^−/−^, n = 9 mice). (**c, d**) T2 weighted MR imaging and cresyl violet staining of brain injury following 24 h reperfusion in C57BL/6 and Cx40^−/−^ mice. (**e**) Summary data and non-parametric analysis of tissue injury in animals subjected to stroke and 24 h reperfusion (T2 weighted imaging: *U* = 19.50, P<0.05, Mann-Whitney *U* test; C57BL/6, n = 9 mice; Cx40^−/−^, n = 9 mice), (cresyl violet: *U* = 18, P<0.05, Mann-Whitney *U* test; C57BL/6, n = 9 mice; Cx40^−/−^, n = 8 mice). Edema volume was not different between the two animal groups. * denotes significant difference between groups.

### Virtual cerebral arterial network predicts the ascension of conducted responses

To assess the impact of conduction on integrated network function, we established and utilized an *in silico* model of vascular communication. Network architecture based on observations from Shih et al. (2009) (24) consisted of a surface vessel plexus, adjoined to six penetrating arterioles (Fig. 4a,b); each composed of smooth muscle and endothelial cells with defined structural and electrical properties. Simulations entailed voltage clamping an endothelial segment in one, two or all penetrating arterioles while monitoring responses throughout the entire network structure (Fig. 4c,d). As the number of stimulated penetrating arterioles increased, the electrical (Fig. 4c,e) and vasomotor (Fig. 4f and “SI Appendix, Fig. 9”) responses progressively ascended into the surface vessel network, where their impact on blood flow delivery would be greater. The ascension process was diminished but remained intact when the voltage clamp was reduced from −55 to −45 mV (“SI Appendix, Fig. 10”). Simulating the increase in endothelial-to-endothelial coupling resistance in Cx40^−/−^ arteries, impaired the ascension of electrical/vasomotor responses (Fig. 4c,e,f and “SI Appendix, Figs. 11,12”). This attenuation was observed irrespective of whether the distal arteriolar segments were hyperpolarized or depolarized, as quantitated in smooth muscle cells located at one site in the penetrating arteriole and at two sites in the cortical surface vessels (Fig. 4d-f and “SI Appendix, Figs. 9-12”). Furthermore, our *in silico* network simulations also equated varying endothelial-to-endothelial coupling resistance to direct changes in ascension of electrical and vasomotor responses (“SI Appendix, Figs. 11,12”).

### Conducted responses are integral to neurovascular coupling function

Past studies of functional hyperemia have implied that conduction plays a central role in coupling blood flow delivery with energetic demand (25-28). To rigorously test this perspective and to place it within our structured biophysical framework, mice were implanted with cranial windows, and trained to run on an exercise ball with their head fixed to an external apparatus. The cerebral vasculature was then labeled with fluorescein isothiocyanate and imaged by 2-photon microscopy while the barrel cortex was activated by whisker stimulation (Fig. 5a-c). The resulting dilatory response was monitored every 300 µm, starting in a penetrating arteriole (upstream of the barrel cortex) and into the adjoining cortical surface artery (Fig. 5c,d). Whisker stimulation in C57BL/6 mice initiated a marked dilation of penetrating arterioles that ascended upstream, with its magnitude decreasing as it advanced along the surface vessel (Fig. 5e,f). The spreading vasomotor response was strikingly reduced in Cx40^−/−^ mice; changes at the 600 or 900 µm sites were minimal compared to controls (Fig. 5e,f). A non-linear regression analysis of the sequential measurement sites revealed that the rate constant of decay, decreased significantly in Cx40^−/−^ mice (λ=501±269) compared to C57BL/6 mice (λ=1385±304; “SI Appendix, Fig. 13”). These findings highlight that conduction is an integral aspect of cerebrovascular control and neurovascular coupling responses to deep brain activity. Note that Cx40 is not a neuronal gap junctional subunit in adult mice and that Cx40^−/−^ animals displayed no change in general behavior.

### Conduction is crucial for blood flow homeostasis during cerebral ischemia

Surface vessel dilation occurs under pathophysiological conditions such as stroke to reroute blood flow to penumbral tissue. This response is observable in humans and animals alike and the redirection of blood notably improves recovery and tissue salvage (29). We hypothesized that vascular conduction is key to surface vessel dilation and consequently, a strategy was implemented to test this concept. Focal cerebral ischemia was induced by occluding the middle cerebral artery and change in perfusion was monitored using Arterial spin labelling-magnetic resonance imaging (ASL-MRI) in C57BL/6 and Cx40^−/−^ mice. Blood flow changes were quantified in two cortical and two striatal regions; Cortical/striatal region 1 demarks the watershed region supplied by middle/anterior or middle/posterior cerebral arteries while cortical/striatal region 2 represents the area supplied by the middle cerebral artery (Fig. 6a). In sham controls, cortical/striatal blood flow averaged between 175-195 ml/100g/min and was similar in C57BL/6 and Cx40^−/−^ mice (Fig. 6b). Occluding the middle cerebral artery in C57BL/6 mice starkly reduced cortical and striatal blood flow (Fig 6c,d) and a partial but robust recovery was observed during short term reperfusion (3h) (Fig. 6c,d). In comparison, the blood flow response to stroke was notably lower in Cx40^−/−^ mice, especially in the cortical/striatal watershed regions, achieving statistical significance only in cortical region 1 (Fig. 6c,d). Similarly, flow responses during reperfusion were also compromised in Cx40^−/−^ animals, notably in the cortical regions and in watershed striatal region 1 (Fig. 6c,d). Diminished blood flow delivery in Cx40^−/−^ mice cannot be explained by anatomical differences in cerebral vasculature or altered hemodynamics. Indeed, Cx40^−/−^ animals are shown to have a similar number of pial surface vessels (30) and our analysis revealed identical heart rates (C57BL/6; 608±97 vs Cx40^−/−^; 609±84), in comparison to controls. Mean arterial blood pressure was modestly higher in Cx40^−/−^ mice (135.2+4.5 vs 115.7±3.1; U = 14, P<0.05, Mann-Whitney *U* test; control, n=10 animals; Cx40^−/−^, n=10 animals), a physiological variable otherwise linked to better perfusion responses during and after stroke (31, 32).

Stroke injury evolves during the acute phase and if perfusion is sustained, brain injury will be moderated. To test the linkage between conduction, blood flow and stroke injury, the reperfusion period was extended to 24 hours in a subset of animals, and measures of perfusion were complemented with assessments of tissue injury. Consistent with our previous observations, brain perfusion was decidedly lower in Cx40^−/−^ mice compared to C57BL/6 controls. This difference persisted after 24 hours of reperfusion, particularly in the cortical regions and in watershed striatal region 1 (Fig. 7a,b). Blood flow in striatal region 2, a region fed by the middle cerebral artery and encompassing the core of ischemic damage, was similar in C57BL/6 and Cx40^−/−^ mice during stroke and early reperfusion although there was a trend of flow recovery in the C57BL/6 mice after 24 hours of reperfusion compared to Cx40^−/−^ animals (Fig. 7a,b). The marked reduction in perfusion translated to augmented tissue injury in the Cx40^−/−^ animals, as MR and histological examinations confirmed an increase in stroke volume (Fig. 7c-e). Interestingly, ischemic damage was similar in the core MCA territory but showed augmentation in adjacent brain levels, indicative of greater penumbral tissue loss in Cx40^−/−^ mice (Fig. 7d). Collectively, these data confirm the decisive role for vascular conduction in vessel responses to stroke and in the preservation of salvageable brain tissue.

## Discussion

Matching blood flow delivery to neural activity requires the cerebral arterial network to respond in an integrated manner (1, 2). The coupling process begins with a neural-derived stimulus and a focal vasomotor response, which must subsequently spread along arteries, to induce the broader changes in resistance needed to impact blood flow magnitude. Spreading vasomotor responses are largely independent of metabolite diffusion or pressure/flow change (7) but require the sharing of a common signal among neighbouring vascular cells (4, 5). This study illustrates that charge is that signal and its conduction, first along the endothelium and then to the overlying smooth muscle, is the mechanism that coordinates contractile activity. This perspective is grounded in the knowledge that cerebrovascular cells are interconnected, express gap junctional Cx subunits (14-16), and that vasomotor responses can robustly spread along, both human and mice cerebral arteries. Building on this foundation, we initiated a quantitative *in silico* analysis to show how the structural/electrical properties of vascular cells are ideally designed to enable the ascension of vasomotor responses from penetrating arterioles to the surface vessel network. Vasomotor ascension was observed in awake mice subjected to barrel cortex stimulation, a response markedly compromised in mice lacking Cx40, a key gap junctional subunit between neighbouring endothelial cells. Impairing the ability of electrical/vasomotor responses to ascend into surface arteries also weakened flow regulation during/following stroke, promoting brain injury. We conclude that conducted responses are an integral part of the coupling mechanisms, needed to optimize the delivery of oxygenated blood to active neural cells, under a range of functional paradigms.

### Conduction is a functional necessity and is enabled by structural connectivity

The study of neurovascular coupling is often reduced to identifying stimuli and the cells from which they are released. This focus routinely overlooks significant conceptual issues, the most important being that vascular resistance is broadly distributed within an arterial tree (2). As such, stimuli that are discretely released and of limited diffusional range cannot mobilize a sufficient proportion of the vascular structure to augment perfusion magnitude (5-7). The vasculature must therefore be encoded with a mechanism to transcribe local neural activity and the stimuli they produce into coordinated multi-segmental responses. While vascular coordination has been traditionally attributed to presumptive changes in pressure and flow, the shortcomings of this logic are clear: upstream change in hemodynamics due to focal stimulation do not need to be substantive or sustained (33). It is within this context that intercellular signalling is considered, with gap junctions enabling electrical, and their corresponding vasomotor responses to conduct along arteries and branch points.

For electrical responses to conduct along cerebral arteries, a defined structural arrangement must exist to facilitate signal exchange among neighboring cells. Using longitudinally sectioned mouse cerebral arteries and electron microscopy, this study documented sites of contact between endothelial cells, between smooth muscle cells, and between the two cell layers (Fig. 1). Endothelial-to-endothelial contacts were particularly abundant with electron dense pentalaminar membranes, an observation indicative of robust gap junctional protein expression. In contrast, sites of contact between smooth muscle cells and the two cell layers were modest in number, with the latter typically observed as a projection extending through “holes” in the internal elastic lamina layer (18-19). We subsequently observed Cx40 expression in the endothelial layer, most notably at homologous contact sites, consistent with their role in charge conduction. Cx37 and Cx43 were also expressed at endothelial cell boarders and near IEL holes (“SI Appendix, Fig. 2”). Translational investigations detected varying levels of Cx37, Cx40, Cx43 and Cx45 mRNA in human cerebral arteries and confirmed the presence of Cx40 protein in the endothelial cell layer (“SI Appendix, Figs. 1, 6”). Overall, our observations align well with findings from the peripheral vasculature and are important in two aspects (18-20). First, they support the postulation that coupling resistance is markedly lower between endothelial cells compared to other contact sites and secondly, that Cx40 deletion will preferentially increase endothelial-to-endothelial coupling resistance.

### Endothelial gap junctions facilitate conduction of vasomotor responses

Building upon structural observations, e*x-vivo* experiments subsequently provided clear evidence that discrete electrical events, either depolarizing or hyperpolarizing in nature, can elicit conducting vasomotor responses in cerebral vasculature. In detail, we observed in both human and mouse cerebral arteries that the discrete application of ‘high pipette [K^+^]’ stimulus elicited a localized depolarization and constriction that conducted along the entire length of the cannulated artery (Fig. 2). A ‘high pipette [K^+^]’ stimulus consistently initiates robust conduction as it shifts E_K_ strongly rightward (∼15-20 mV depolarization). A ‘low pipette [K^+^]’ stimulus and acetylcholine both induced a comparably small but consistent conducted vasodilation that decayed along surface vessels and penetrating arterioles, both shown to express endothelial Cx40 (Fig. 1 & 3; “SI Appendix, Fig. 1”). While small in magnitude, this response aligns with biophysical expectations. In a confined structure of coupled cells, subtle changes in resting K^+^ channel activity won’t be able to generate sufficient current to initiate a robust endothelial V_M_ response. Robust responses could be initiated if an agent were to: 1) profoundly activate a silent population of K^+^ channels (e.g. small/intermediate conductance Ca^2+^ activated K^+^ channels); or 2) stimulate an enhanced number of endothelial cells. The latter concept is intriguing when considered in the context of stimuli diffusing through interconnected structures (e.g. capillaries).

Similar to vessels from peripheral circulatory beds (4, 20), the endothelium was critical for the longitudinal charge spread, as its elimination prevented vasomotor responses from conducting beyond the point of agent application. The genetic deletion of Cx40, a Cx subunit expressed exclusively in the endothelium, impaired but did not abolish conduction; endothelial Cx37 and Cx43 were also downregulated in these mice (“SI Appendix, Fig. 7”); an effect that could also contribute to the impairment in conduction. This deletion-induced impairment did not affect vessel reactivity to a global K^+^ challenge nor did influence myogenic reactivity of cerebral arteries (“SI Appendix, Fig. 8”). Consecutively, V_M_ arterial measurements confirmed the attenuation of charge spread along the artery in the absence of Cx40^−/−^ in the endothelium. Changes in endothelial-to-endothelial coupling resistance were subsequently calculated using a validated model of electrical communication, modified to convert diameter responses into voltage changes (“SI Appendix, Fig. 3 & 4”). Best fit analysis yielded endothelial-to-endothelial coupling resistances of 1.7 MΩ and 6 MΩ in C57BL/6 and Cx40^−/−^ mice, respectively. The former aligns surprisingly well with direct electrical measurements performed on mice microvascular endothelial cells (34).

### Intercellular signalling drives ascending neurovascular responses

The preceding experiments provided essential knowledge needed to advance a deeper understanding of conduction in integrated network function. In this regard, gap junctional resistance values were incorporated into a virtual network that consisted of a previously described surface vessel network with six adjoining penetrating arterioles (24). With endothelial-to-endothelial coupling resistance set at 1.7 MΩ (C57BL/6, control mice), simulations revealed that stimulating one endothelial segment in a single penetrating arteriole was sufficient to initiate changes in V_M_ and vasomotor tone along the arteriole and marginally in the cortical surface vessels (Fig. 4; “SI Appendix, Fig. 9”). The robustness of this ascending response, albeit hyperpolarization or depolarization, increased substantially as the number of stimulated penetrating arterioles were increased. A similar pattern of ascension was also observed when the strength of the initial hyperpolarizing stimulus was decreased (“SI Appendix, Fig. 10”). This grading of upstream vasomotor behavior is biologically important; as in its absence blood flow delivery cannot be properly matched with varying neural activity. Elevating endothelial-to-endothelial coupling resistance to 6 MΩ (Cx40^−/−^ mice) markedly attenuated the ability of electrical/vasomotor responses to ascend through the arterial tree. These findings establish the importance of endothelial-to-endothelial cell coupling in vascular conduction and illustrates, similar to functional studies (25-28), that compromised coupling would decisively impact functional hyperemic responses. In fact, by varying endothelial-to-endothelial cell coupling, in silico network also indicated its direct correlation to ascending electrical and vasomotor responses (“SI Appendix, Figs. 11 & 12”). We examined this idea experimentally by implanting cranial windows in awake mice and monitoring vasomotor activity prior to and following barrel cortex stimulation (Fig. 5). Notably in control mice, whisker stimulation elicited a dilation that ascended in an electronic fashion from penetrating arterioles to pial arteries. Consistent with conduction driving this upstream response in an endothelial/gap junctional dependent manner, the deletion of Cx40^−/−^ attenuated this functional hyperemic response. These findings confirm that neurovascular coupling is a two-step process, consisting of a stimulus initially driving a focal electrical/vasomotor response, followed by charge spread via gap junctions, altering contractile activity along an extended portion of the arterial network. As expected, Cx40^−/−^ mice display no change in general behavior as Cx40 is not a neuronal gap junctional subunit.

### Conducted response is vital to perfusion homeostasis during stroke

The necessity of conduction to maintain vasomotor activity and perfusion is a new, unexplored concept in the cerebral circulation. While logically framed in the context of functional hyperemia, conduction may also be important in pathophysiological scenarios. Case-in-point is stroke where occlusion of a major cerebral artery dilates surface vessels, rerouting blood from other branches to the tissue at risk (25). This rerouting of perfusion is observable in humans and rodents alike and limits tissue injury, particularly in penumbral (salvageable) regions outside the ischemic core (25, 31). The mechanism underlying surface vessel dilation is unknown, but it follows that conduction, given the external positioning of surface vessels, is a likely player. We tested this hypothesis by subjecting mice to middle cerebral arterial occlusion and monitoring perfusion during stroke and reperfusion (Figs. 6 & 7). Compared to C57BL/6 controls, perfusion was impaired in Cx40^−/−^ mice during stroke and during the early reperfusion phase. This diminution was particularly evident in the cortex and was observed despite a modest rise in blood pressure, predisposing Cx40^−/−^ mice to enhanced blood flow during stroke. As stroke injury development likely depends on sustained blood flow regulation in the surface vessel network (35), we subsequently extended the reperfusion period and paired perfusion data with measurements of tissue injury. The attenuated blood flow responses in Cx40^−/−^ mice during the short reperfusion period were consistent, even after 24 hours of reperfusion and even became more pronounced. These differences correlated positively with augmented tissue injury in Cx40^−/−^ mice, as revealed by T2 weighted imaging and cresyl violet histology. Our findings establish the importance of intercellular signalling and vascular conduction for the maintenance of perfusion during stroke, where the brain is critically vulnerable to injury.

### Summary

The optimization of neural function requires cerebral arterial networks to tune how much and where blood flow is distributed within the brain (1,2). In this regard, investigative efforts have largely focused on identifying key vasoactive agents (e.g. prostaglandins, epoxyeicosatrienoic acids, nitric oxide, adenosine, K^+^ etc.) and the cellular structures from which they originate (2). Deeper discussions of biophysical constraints have been lacking in these examinations, the most important being that focally derived vasoactive agents have a limited diffusional range. Thus, they alone cannot alter tone in a sufficient portion of the arterial tree to effect a sizable change in blood flow magnitude. Our study illustrates that this constraint is overcome in the cerebral arterial networks where focal responses can conduct along arteries and across branch points. The conduction process is facilitated by gap junctions and as our quantitative work show, impairing communication diminishes perfusion and vasomotor control over a range of metabolic challenges. Without electrical signalling to drive conducted vasomotor responses, blood flow delivery cannot be effectively matched to metabolic demand, depleting brain’s energy sources and compromising its function.

## Materials and Methods

### Animals and reagents

All procedures were approved by the Animal Care Committees at the Universities of Calgary and Western Ontario and complied with the Canadian Council on Animal Care and ARRIVE (Animal Research: Reporting *In Vivo* Experiments) guidelines. All animals (See “SI Appendix, Table 1” for details) were male, 8-10 weeks of age and within 20-25 grams body weight; were group housed with a 12-hour light/dark cycle and with free access to food and water. Further details could be found in the “SI Appendix (text)”. Experimental sample (n) numbers can be found in “SI Appendix, Tables 1 & 2”; all reagents were purchased from Sigma-Aldrich (MO, USA), unless otherwise stated.

### Human tissue experiments

A total of 11 human cerebral arteries (3 for conduction assay; 2 for immunohistochemistry; 6 for qPCR) were collected from 6 patients (3 males/3 females), (27-59 years of age) undergoing epileptic resection surgery. All experiments involving human tissue was approved by the Health Sciences Research Ethics Board at the University of Western Ontario (HSREB ID#107196). All samples were obtained after written, informed consent from patients. Resected brain sections were collected in a sealed container with saline solution, on ice and was transported to the lab in an insulated carrier bag. Freshly isolated cerebral arteries were used for conduction assay while isolated arteries were frozen in liquid nitrogen and stored at −80 for qPCR experiments.

### Myography, conduction assay and arterial membrane potential

Animals were euthanized by carbon dioxide asphyxiation and brains were carefully removed and placed in phosphate-buffered saline solution (pH 7.4) containing (in mM) 138 NaCl, 3 KCl, 10 Na_2_HPO_4_, 2 NaH_2_PO_4_, 5 glucose, 0.1 CaCl_2_, and 0.1 MgSO_4_ (36-38). First and second order middle cerebral arteries were dissected away from surrounding tissue and cut into 2.5 mm segments. The segments were then cannulated in a customized arteriograph and superfused with warm (37^°^C) physiological salt solution (PSS; pH 7.4) containing (in mM) 119 NaCl, 4.7 KCl, 1.7 KH_2_PO_4_, 1.2 MgSO_4_, 1.6 CaCl_2_, 10 glucose, and 20 NaHCO_3_. Arterial diameter was monitored using an automated edge detection system (IonOptix, MA, USA) coupled to a 10X objective. In a subset of experiments, a middle cerebral artery with penetrating arteriole was isolated, triple cannulated and pressurized (50 mmHg) so that conduction experiments could be performed on the penetrating arteriole. Please see “SI Appendix, (text)” for more details.

The conducted response was assessed by placing a narrow bore (1-2 μm diameter) glass micropipette backfilled with KCl (500 mM) near one end of a pressurized human/mouse cerebral arteries (37, 38). Using pressure ejection (30 psi, 10s pulse), KCl was applied to a small portion of the artery inducing depolarization and a local constriction (25-30 µm) that represents ∼50% the maximal constrictor capacity; within this range changes in membrane potential lead to a graded vasomotor response (37). Our approach yields a 15-20mV change of V_M_ at the stimulation site (Figure 2e); a response that corresponds with a ∼50 mM rise in local [K^+^] (23). Arterial diameter was monitored at sites 0, 450, 900, 1350 and 1800 µm distal to the point of agent application. Conducted dilatory responses were assessed after pressure injection of acetylcholine (1 mM) or K^+^ (25 mM); arterial diameter changes were monitored at the 0 and 450 µm sites on an isolated surface vessel or a penetrating arteriole coupled to a surface vessel (triple cannulation preparation). We chose to use small-bore pipettes backfilled with high stimulant concentrations as this approach minimizes the volume of fluid ejected onto the preparation. Superfusate flow then dilutes and crucially, keeps stimulants away from the conducted sites, positioned upstream from the site of agent application. Vasomotor responses were calculated as the difference in vessel diameter before and after stimulation with agonist. The role of the endothelium was investigated by comparing the conducted vasomotor responses prior to and following its removal, by passing an air bubble down the vessel lumen. Each group consisted of 7-10 vessels, each from a different animal (see “SI Appendix, Table 1” for details).

Arterial membrane potential (V_M_) was monitored with a glass microelectrode backfilled with 1M KCl (tip resistance = 75-100 MΩ) inserted into the vessel wall. Cerebral arteries were pressurized to 50 mmHg and V_M_ was assessed in the presence of nifedipine (200 nM) to minimise changes in vasomotor tone. A successful impalement consisted of: 1) a sharp negative deflection in V_M_ upon insertion; 2) a stable recording for at least one minute; and 3) a sharp return to baseline upon retraction.

### Quantitative polymerase chain reaction (qPCR)

Cerebral arteries were collected from resected brain tissue (6 arteries/4 patients) and middle cerebral arteries (MCA) were collected from C57BL/6 and Cx40^−/−^ animals (6-7 animals per group, details in “SI Appendix, Table 1”). Total RNA was isolated using RNeasy plus micro kit (Qiagen, ON, Canada) and reverse transcription was performed with the Quantinova reverse transcription kit (Qiagen); reaction mixtures lacking reverse transcriptase was used as negative control. Following standardization to the reference genes, glyceraldehyde 3-phosphate dehydrogenase (GAPDH) (human) or α-actin (mouse); test genes (vascular connexins) in human vessels were normalized relative to human whole heart (RNA purchased from Clontech) and in mouse, were expressed relative to C57 (Cx40^+/+^) vessels, using the relative expression software tool (REST) (39). Please see “SI Appendix, (text)” for more details.

### Electron microscopy and connexin immunohistochemistry

For electron microscopic examinations, animals were perfusion fixed in 2% paraformaldehyde, 3% glutaraldehyde in 0.1 M sodium cacodylate, with 10 mM betaine and 0.1 M sucrose. Vessel segments, ∼3-5 mm long, were embedded in Araldite 502 using standard procedures and sectioned longitudinally. Sections were imaged at 4 MP on a transmission electron microscope (JEOL (Australasia) Pty. Ltd.).

For Cx immunohistochemistry, animals were perfusion fixed in 2% paraformaldehyde in 0.1 M PBS. Middle cerebral arteries were dissected, opened and pinned (en face preparation) on Sylgard (WPI, USA); penetrating arterioles were also dissected and mounted. Sections were compared against appropriate controls (without primary antibody, Cx40 binding peptide). Propidium iodide or DAPI were used to label nuclei as needed. Please see “SI Appendix, (text)” for details; a list of primary/secondary antibodies used can be found in “SI Appendix, Table 4”).

### Computational modeling

A single vessel and an arterial network model were constructed using a generative modeling approach (36). A single vessel model of defined length and diameter was comprised of one layer of endothelial and two layers of smooth muscle; each was represented 3D space with cells of a particular volume. Homocellular gap junction couplings are defined using a pattern generator that specify interacting cells (a principle that only specifies links among model objects and not specific cell coordinates). Each endothelial and smooth muscle cell was given 6 homocellular couplings and conductance was set according to past calculations (3, 40). Ionic conductance was represented by polynomial fits of whole cell IV curves from cerebral arterial endothelial and smooth muscle cells (“SI Appendix, Fig. 3”). While IV curves are generally linear over the physiological V_M_ range, a number of the underlying channels do display voltage dependent properties which could impact conduction (38). Arterial V_M_ was subsequently coupled to diameter through a sigmoid function (“SI Appendix, Fig. 4”). Using a best fit approach, this computational model was used to determine the endothelial-to-endothelial coupling conductance from data collected in Figure 2. Please see “SI Appendix, (text)” for further details.

### Neurovascular coupling experiments and Two-photon microscopy

Animal surgery and training were performed as described previously (41). Briefly, mice were anesthetized with isoflurane (5% for induction and 1-1.5% for maintenance), mounted on a stereotaxic frame and body temperature maintained using a feedback control system for all surgeries. One week before imaging, a head bar was implanted over the interparietal bone and glued in place with cyanoacrylate without creating a cranial window. An observation well was progressively built around the future cranial window using dental cement, for a water dipping objective. After two days of rest, animals were given two training sessions (30-45 mins) on a spherical air-supported treadmill. After at least two more rest days, craniotomy was performed to generate a cranial window (3×3 mm square) over the barrel cortex. Animals were then positioned on the spherical treadmill with the head immobilised using the head bar and activity was constantly monitored using a video camera. FITC-dextran was injected through a tail artery cannula to label the vasculature. Imaging of the microvascular network was performed when the animals displayed signs of alertness (vigorous running) and had passive whisker movements. A 5 sec air puff was delivered to the contralateral side to deflect all the whiskers while vascular responses were imaged using a custom built *in vivo* two-photon microscope equipped with a Ti:sapphire laser (Coherent Chameleon, Ultra II, ∼4 W, 670-1080 nm, ∼80 MHz, 140 fs pulse width) and controlled by Scan Image software. Please see “SI Appendix, (text)” for further details.

## Mouse model of ischemic stroke

Animals were anesthetized using 1.5% isoflurane (30% O_2_, remainder N_2_O) and body temperature was maintained using a feedback-controlled heating system. Focal cerebral ischemia was induced using an intraluminal technique (42). Briefly, a midline neck incision was made and the left common and external carotid arteries were isolated and ligated. A microvascular clip was temporarily placed on the internal carotid artery and an 8-0 silicon-coated nylon monofilament (Doccol, USA) was directed through the internal carotid artery until the origin of MCA. After induction of ischemia, animals were transferred to the MRI system for imaging. Following 60 minutes of ischemia, the monofilament was withdrawn, and reperfusion was monitored at 1, 2 and 3 hours. In a subset of animals, stroke was induced/monitored, and reperfusion was extended to 24 hours during which a repeat scan was performed. Sham controls underwent anesthesia and midline incision, without manipulation of the arteries. Anesthesia was maintained by monitoring the heart rate and respiration, throughout the experimental procedure.

### Magnetic resonance imaging and analysis

MR imaging and analysis were performed on a 9.4T/21 cm horizontal bore magnet (Magnex, UK), a Bruker BGA-12S gradient system and an Avance II hardware (Bruker, Germany) as described (43). The animal’s head was secured in a 35mm quadrature volume radiofrequency coil, positioned at the centre of the magnet. While inside the magnet, animal anesthesia and body temperature were monitored and maintained. Each scanning session acquired a set of T_2_ weighted (T_2_W) images and a Perfusion-Weighted (PW) map, for investigating the changes in tissue injury and cerebral blood flow, respectively (44, 45). T_1_ Weighted (T_1_W) maps were also acquired from corresponding slice location as PW scans. Details of scan parameters are provided in the “SI Appendix, (text)”.

Cerebral perfusion was calculated in a total of 13 Regions of Interests (ROI’s) as:

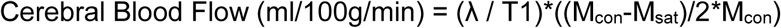

where T1 was calculated from T1 weighted images and partition coefficient of water (λ) was assumed to be 0.9 (46). M_con_ is the intensity of control images and M_sat_ is the intensity of the images acquired after arterial inversion (42). Detailed analysis parameters are provided in the “SI Appendix, (text)”.

### Histology

After MR imaging (24 h reperfusion endpoint), animals were euthanized and transcardially perfused, first with heparinized saline and then with normal saline (47). Brains were removed, frozen on dry ice and cut into 20 µm coronal sections using a cryostat. Brain sections 1 mm apart were stained with cresyl violet and image analysis software (ImageJ; NIH) was used to outline the border between infarcted and healthy tissue (42). Infarcted area was quantified by subtracting the area of non-lesioned ipsilateral hemisphere from that of the contralateral hemisphere. Infarct volume was calculated by integrating infarcted areas from 6 equally spaced rostrocaudal sections. Edema or brain swelling was calculated as absolute difference between ischemic and non-ischemic hemispheres (42, 47, 48). An examiner blinded to tissue identity performed the analysis.

### Blood pressure and heart rate assessment

The non-invasive assessment of blood pressure was performed on conscious C57BL/6 and Cx40^−/−^ animals (10 animals per group) using the non-invasive CODA tail-cuff system (Kent Scientific, CT, USA) as previously described (49). To minimize anxiety, animals were properly acclimatized, and body temperature was maintained with a warming platform. Mean arterial blood pressure and heart rate was averaged from two experiments, each consisting of up to 15 individual trials.

### Statistical analysis

Data sets are expressed as box and whisker plots and P < 0.05 was considered statistically significant. Power analysis was done to calculate sample size sufficient for obtaining statistical significance and n numbers reported in previous publications were considered, in experiments involving vessels and animals. Significant results are indicated in the graphs after animal groups were evaluated and compared for differences using non-parametric Mann-Whitney U test (single tailed) (GraphPad PRISM 4.0, CA, USA), performed at individual time points. To assess differences in conduction decay *in vivo*, individual data sets were fitted to an exponential function f(x) = k*exp(−x/lambda), using the nonlinear least-squares Marquardt-Levenberg algorithm (Gnuplot version 5.2). A single tailed-t-test compare the summative decay constant from wild type and Cx40^−/−^ mice. P < 0.05 was considered statistically significant. Experimental design and data presentation are reported in compliance with the ARRIVE guidelines.

## Acknowledgements

This work was supported by operating grants from the Canadian Institute of Health Research (CIHR) to D.G.W (377412) and G.R.J.G (130233). D.G.W is the Rorabeck Chair in Molecular Neuroscience and Vascular Biology at the University of Western Ontario; G.R.J.G is a Canada Research Chair (Tier 2). A.Z was supported by postdoctoral fellowships from the University of Calgary (Eyes High), Alberta Innovates Health Solutions (AIHS) and CIHR. C.H.T-T was supported by a postdoctoral fellowship from AIHS. S.L.S was supported by the Brain Foundation and Diabetes Trust, Australia.

The authors would like to acknowledge the excellent, general technical support of Suzanne Brett (University of Western Ontario); Tadeusz Foniok and David Rushforth (Experimental Imaging Center, University of Calgary) for assistance with MR experiments; Dr. Frank Visser (HBI Molecular Core Facility, University of Calgary) for assistance with qPCR experiments; Dr. Robert Gros (Robarts Research Institute, University of Western Ontario) for assistance in blood pressure/heart rate experiments and Michelle Kim (Robarts Research Institute, University of Western Ontario for general technical assistance. Cx40^−/−^ animals were a kind gift from Dr. Janis M. Burt (University of Arizona).

