## Supplementary Information for "Intercellular Conduction Optimizes Arterial Network Function and Conserves Blood Flow Homeostasis during Cerebrovascular Challenges"

#### Materials and Methods

Detailed descriptions of materials and methods are included under respective sections.

#### Animals and reagents

$Cx40^{-/-}$ mice were obtained from Prof. Janis M. Burt (University of Arizona) and are available through Jackson laboratory (B6.129S4-Gja5tm1Paul/J) (Stock No: 025697). Female $Cx40^{-/-}$ were bred with C57BL/6J (C57BL/6) and $Cx40^{-/-}$ offsprings were identified (genotyped) and backcrossed for >4 generations. C57BL/6 mice were used as controls in all experiments. Animals within strains were randomly assigned into different experimental groups. One mouse (C57BL/6 group) died during stroke/MR imaging in the 3 h reperfusion experiments. Two mice were also lost (one each from C57BL/6 and $Cx40^{-/-}$ groups) during the 24 h reperfusion experiments. One brain could not be collected ($Cx40^{-/-}$ group) for cresyl violet staining, as the animal did not recover after the MR session.

#### Myography, conduction assay and arterial membrane potential

Arteries were equilibrated for 30 mins at 15 mmHg intravascular pressure and contractile responsiveness assessed by brief (~10s) exposure to 60 mM KCl in the bath solution. Following equilibration, intravascular pressure was raised to 50 mmHg (in vivo pressure). Maximal diameter was obtained by superfusing with a Ca2+ free PSS containing 2 mM EGTA. Small arterial branches were carefully closed by pinching with forceps, without damaging the main vessel segment. Small caliber middle cerebral arteries were chosen, as a suitable length of vessel (2000 µm) could be isolated with minimal branching; this avoids (a) strong end effects where electrical events reflect back through the vessel and (b) the inclusion of large charge sinks that compromise charge spread.

#### Quantitative polymerase chain reaction (qPCR)

Real-Time PCR was performed using Prime Time qPCR primers (Integrated DNA Technologies, IA, USA) (Supplementary Table 3), the Kapa SYBR Fast Universal qPCR kit (Kapa Biosystems, MA, USA) and cDNA template from cerebral arteries (1ng of template per reaction). The running protocol employed an Eppendorf Realplex 4 Mastercycler (ON, Canada) and was 45 cycles in length (95˚C for 5 seconds, 55˚C for 15 seconds and 72˚C for 10 seconds). PCR specificity was confirmed by dissociation curve analysis and template controls which yielded no detectable fluorescence. Primer efficiencies were determined experimentally and ranged from (80-100%).

#### Electron microscopy and connexin immunohistochemistry

Arterial segments were washed with PBS and incubated for 2 h at room temperature in a blocking buffer containing 1% BSA and 0.2% Triton-X in PBS. Segments were then incubated overnight with primary antibody against Cx37, Cx40 and Cx43 at 4°C followed by incubation with secondary antibody at room temperature for 2 h. Tissue was then mounted in anti-fade mounting media and examined under a FV1000 confocal microscope (Nikon Instruments Inc.) with uniform settings.

#### Computational modeling

In order to create a 3D network, coordinates of vessel bifurcations and network ends were obtained from Shih et al 2009 (1). Assuming vessels to be straight cylinders, vessel lengths and connectivity were calculated (meandering vessels were approximated using piecewise linear curve fits (2). Each surface vessel was covered with two smooth muscle layers and was assumed to have a resting diameter of 75 µm, whereas penetrating arterioles were 400 µm long, covered with a single layer of smooth muscle and assumed to have a resting diameter of 30 µm. At each bifurcation, endothelial cells at the end of an incoming branch were divided into two sets; the number of ECs in each set being dependent on the ratio of ECs in the two opposing branches.

Endothelial cells within each set were coupled to endothelial cells from a single opposing branch while smooth muscle cells, owing to their circumferential orientation, were coupled to at least one smooth muscle cell from each opposing branch, at the bifurcation (3).

IV-curves of cerebral endothelial and smooth muscle cells were obtained by patch clamping of isolated vascular cells in whole-cell mode, in physiological solutions (4). Polynomial fits (Supplemental Fig. 3) were described by:

$I_{m,EC}(V_M) = 1.97 + 66.5 \cdot 10^{-3} V_M + 562 \cdot 10^{-6} V_M^2 + 3.98 \cdot 10^{-6} V_M^3$

$I_{m,SMC}(V_M) = 2.60 + 0.148 V_M + 2.97 \cdot 10^{-3} V_M^2 + 2.57 \cdot 10^{-5} V_M^3 + 7.55 \cdot 10^{-8} V^4{}_M^4$

A $V_M$,rest of -40 mV was achieved by simple coordinate transformation of the $V_M$-axis. Following simulation of $V_M$ within each cell of the vascular network, the electrical response was translated into a change in inner diameter using the sigmoid function:

$$D(V) = D_{max} \cdot \left(1 - \frac{1}{1 + exp\left(K_m \cdot (-V + V_{mrest})\right)}\right)$$

where $V_M$-rest is resting potential that corresponds to the electrical model. For penetrating arterioles, $D_{max}$ = 50µm and for surface vessels $D_{max}$ = 120µm.

To estimate $K_m$, a constant determining the dynamic range of excitation-contraction coupling, a minimal diameter, $D_{min}$ was added to the sigmoid and this function was fitted to the observed relationship between simultaneously measured $V_M$ and outer vessel diameter in cerebral vessels (see Supplemental Fig. 4); from the fit, $K_m$ = 0.134.

**Neurovascular coupling experiments and Two-photon microscopy**

Prior to surgery, the animals were treated with Dexamethasone 21-phosphate disodium salt (anti-inflammatory drug, 4 mg/ml, Sigma) and Buprenorphine (analgesic). Bone and underlying dura mater were removed. A drop of artificial cerebrospinal fluid (ACSF) was then applied to the surface of the cortex; the open window was covered with a cover glass and sealed using cyanoacrylate and dental cement.

Vasomotor responses of the pial and penetrating arterioles were monitored using xy raster scanning (0.98 to 7.8Hz) while the animals were running. Peak vasomotor response (△D) was calculated as the difference in vessel diameter at rest (2 sec time averaged) and following an air puff stimulation. 3D rendering of the microvascular network was done using the 3D viewer plugin (5).

**Magnetic resonance imaging and analysis**

The T2W scan (Repetition time of 4s) was acquired with a field of view of 2cm2, a 128 X 128 matrix and thirty-two continuous slices (0.8mm thick) with an inter-echo spacing of 10 ms. PW map was obtained in a single slice (1 mm) using a continuous arterial spin labelling technique targeted at the mid-striatum region (anatomical centre of MCA supplied region) as described previously (6, 7). A 2s long radiolabelling pulse was applied to the neck vessels in the presence of a 2G/cm gradient followed with a delay of 400 ms by a HASTE (Half-Fourier Acquisition Single Shot Turbo Spin Echo) sequence using a repetition time of 3000 ms, Effective Echo Time of 13.3 ms and a RARE factor of 36, for a total of 16 averages. Magnetization in unsaturated control images were obtained by using the same radiofrequency excitation applied symmetrically, opposite to the labeling plane, to help eliminate magnetization transfer effects.

Absolute blood flow values obtained as below zero were considered as no flow regions (0 ml/100g/min), in analysis. All MR images were analysed using Paravision 5.1 software and four anatomical (two cortical and two striatal) regions were studied, by forming averages from multiple ROI's. Cortical region-1 represented the watershed region of the cortex supplied by MCA and ACA while cortical region 2 was demarcated in the MCA supplied territory of the cortex and were formed from averages of two and five ROI's, respectively. Similarly, striatal region 1 included the watershed area between MCA and PCA while striatal region 2 was located in the core of MCA supplied territory and was formed from averages of three ROI's, each (Fig. 5a).

**Statistics**

To assess differences in conduction decay between C57BL/6 and Cx40$^{-}$/$^{-}$ mice, conduction data set from each group was fitted to an exponential function f(x) = k*exp(-x/lambda), using nonlinear least-squares Marquardt-Levenberg algorithm (Gnuplot version 5.2). Single tailed- t-test was then used to compare decay constants between groups. $P < 0.05$ was considered statistically significant.

### Supplementary Figures

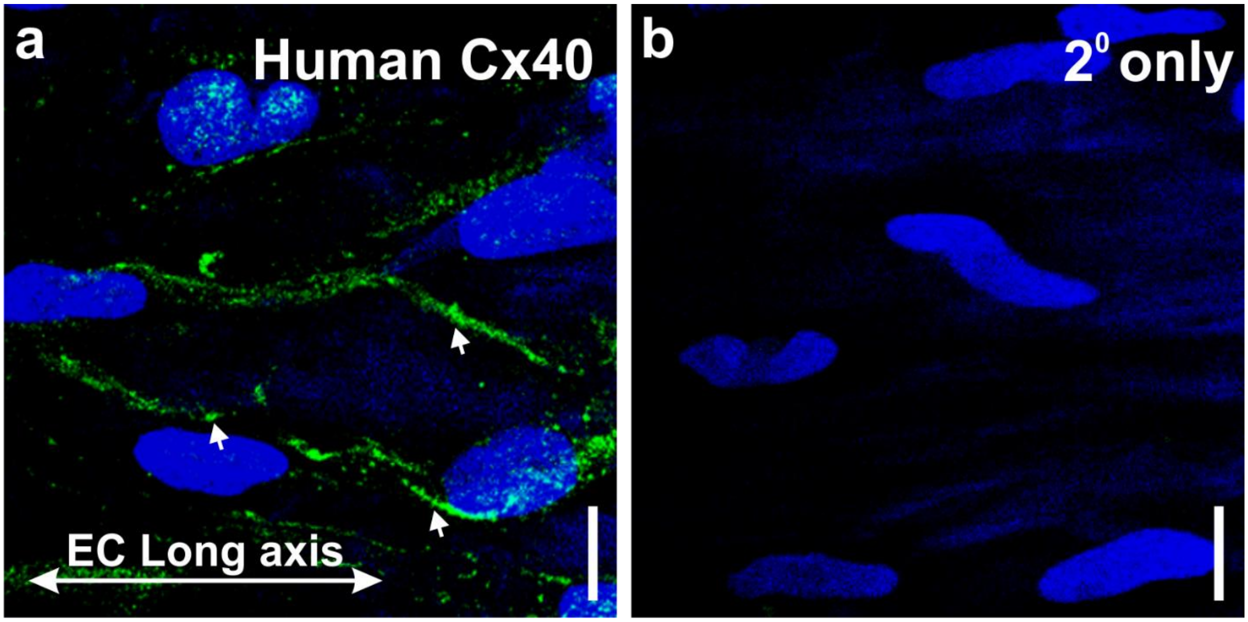


**Figure 1: Endothelial connexin (Cx) 40 expression in human cerebral arteries. (a)** Cx40 (green) was identified in the endothelial layer of human cortical surface artery preparations (en face). Cell nuclei were labeled with DAPI. Arrowheads denote sites on punctate Cx labeling around the periphery of endothelial cells. (**b**) Absence of Cx40 labeling in tissue treated without primary antibody. Scale bar = 10 µm.

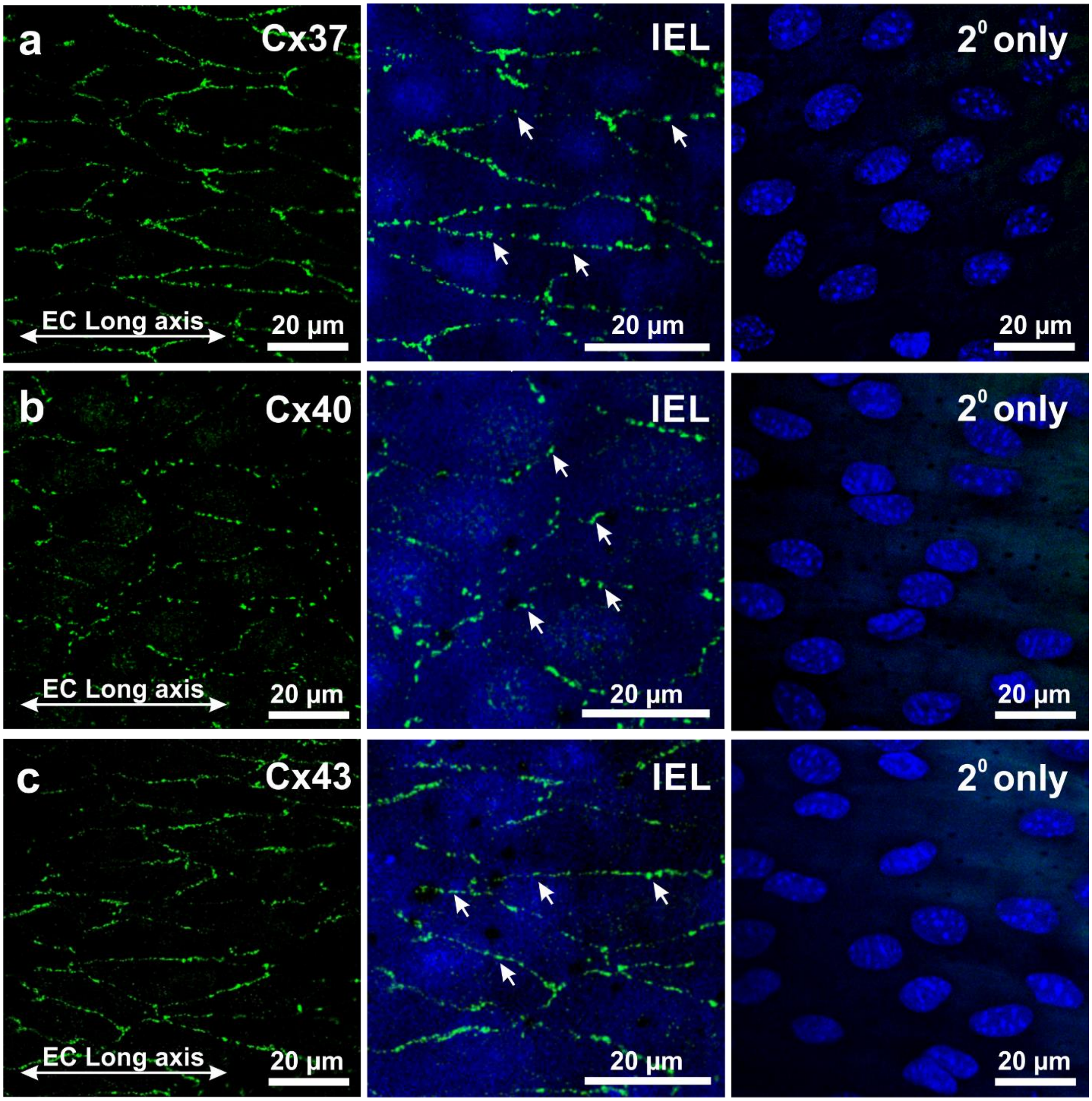


**Figure 2: Endothelial connexin (Cx) are robustly expressed in cerebral arteries from C57BL/6 mice.** **(Left)** Connexin isoforms Cx37 **(a)**, Cx40 **(b)** and Cx43 **(c)** (green), were identified by immunohistochemistry in the endothelial layer of middle cerebral arteries from C57BL/6 mice. **(Middle)** Arrowheads denote sites on punctate Cx labeling around the periphery of endothelial cells and IEL was delineated by autofluorescence (488 nm). (**Right)** Absence of endothelial Cx labeling in opened arteries incubated without the primary antibody; nuclei were labeled with DAPI. Scale bar = 20 µm.

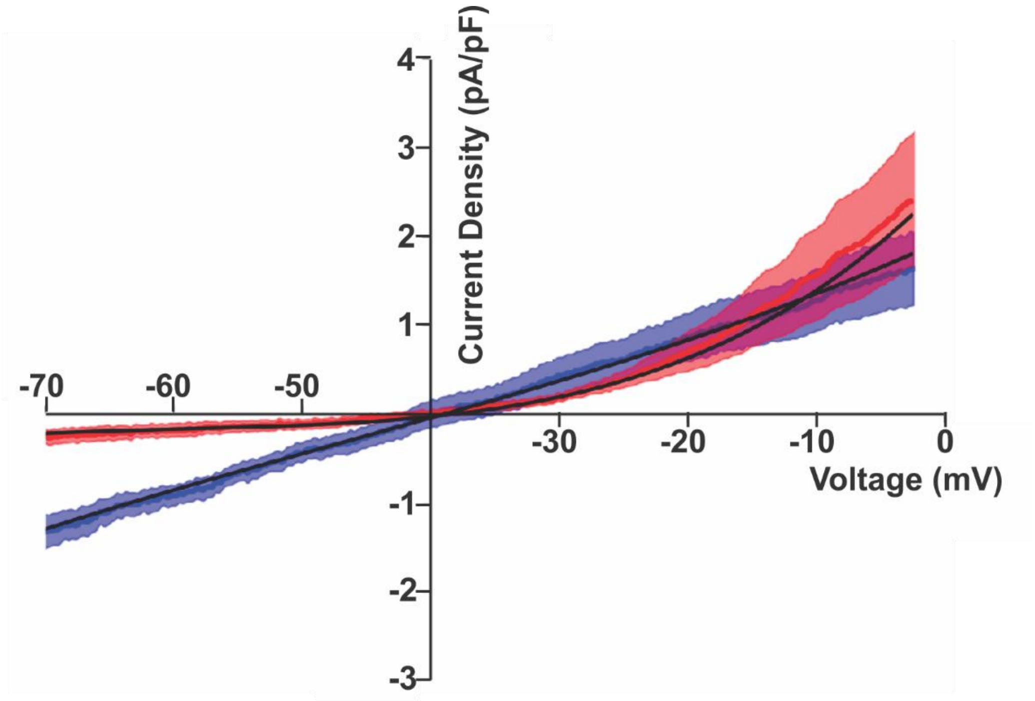


**Figure 3. Polynomial fits of IV-curves from isolated cerebral arterial cells.** Representative whole cell currents from mouse endothelial and smooth muscle cells were collected in physiological bath and pipette solutions (4). Data was fitted with a polynomial curve and values were incorporated into the virtual arterial model to account for ion channel activity. Whole cell capacitance was 8.0 pF and 18.7 pF for the endothelial (blue) and smooth muscle (red) cell, respectively.

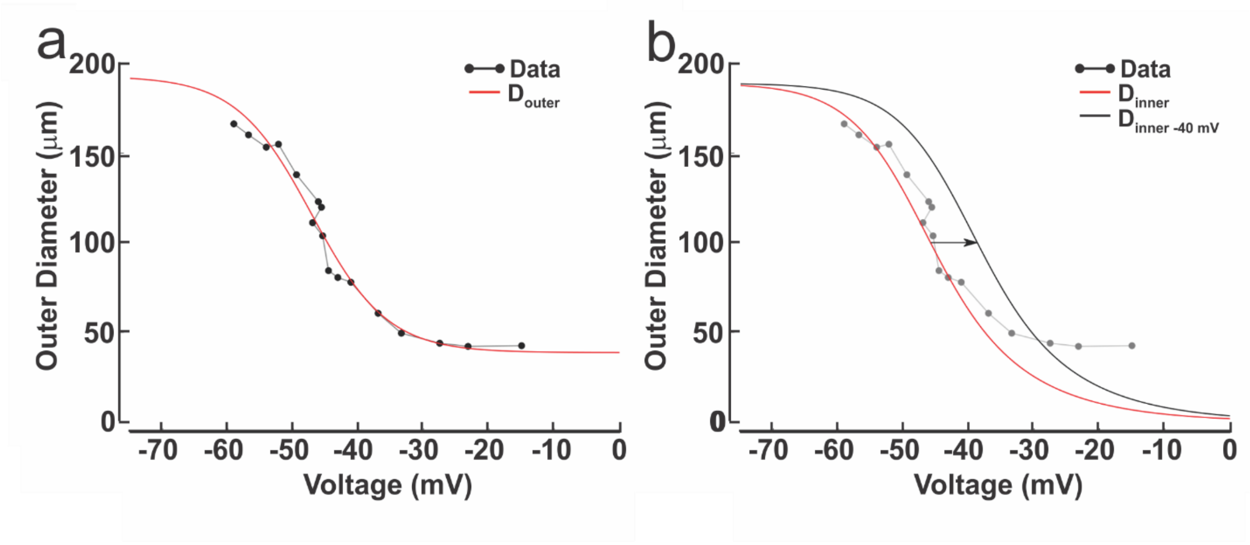


**Figure 4. Calculated relationship between diameter and arterial $V_m$. (a)** Data illustrative of the relationship between arterial $V_M$ and outer cerebral arterial diameter was plotted (2) and a sigmoid function (red curve; see Eq.1) applied. **(b)** Inner diameter was represented by assuming that a vessel is fully contracted at 0 mV and that the cross-sectional vessel area is constant across measurements (red curve). This curve was right-shifted to correspond to a resting $V_m$ of −40 mV; black curve). While either inner or outer diameter could be modelled, the latter was chosen to facilitate comparisons with the neurovascular coupling experiments in Fig. 5.

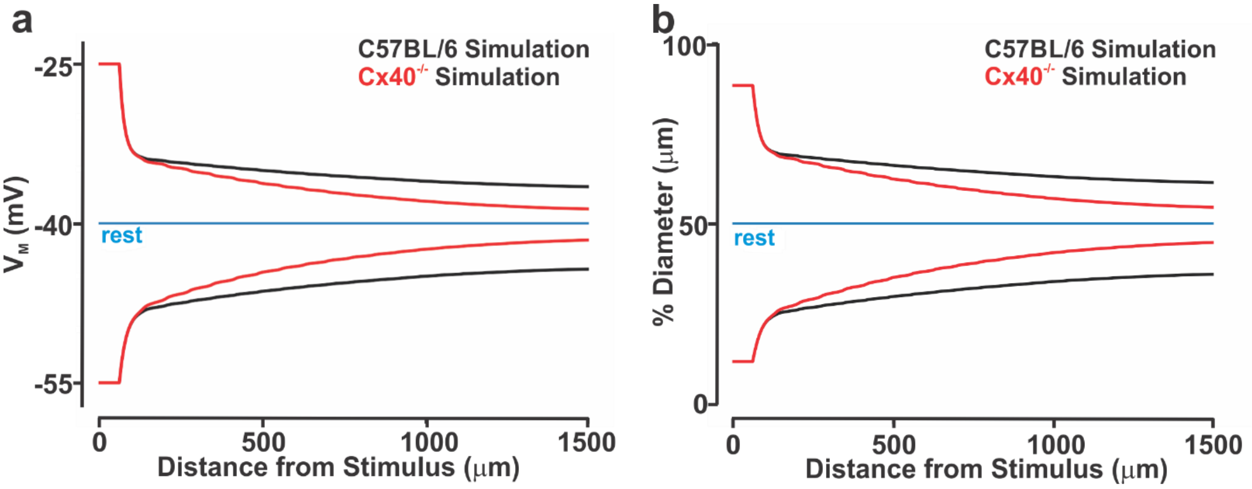


**Figure 5. *In silico* modeling: Electrical communication in a virtual cerebral artery.** A virtual model akin to an isolated cerebral artery (length, 1500μm; diameter, 75 μm; one layer of endothelium and one layer of smooth muscle) was constructed to study electrical communication. **(a, b)** One distal arterial segment was voltage clamped (15 mV negative or positive to resting $V_m$; 250 ms) and the spreading electrical/vasomotor responses were quantitated along the virtual vessel; endothelial-to-endothelial coupling resistance was set to 1.7 (C57BL/6 control) or 6.0 ($Cx40^{-/-}$) MΩ (calculated in Figure 2). Clamping both cell layers elicits a biphasic response due to 1) a strong but local $V_M$ change in smooth muscle, due to its high compound resistance; and 2) a decidedly lower $V_M$ perturbation in the endothelium as charge readily flows along this cell layer. When translated to diameter, this biphasic response is modestly dampened by the sigmoid shape of the $V_M$-diameter relationship. Increasing endothelial-to-endothelial coupling resistance ($Cx40^{-/-}$ simulation) enhanced electrical/vasomotor decay relative to control (C57BL/6 simulation).

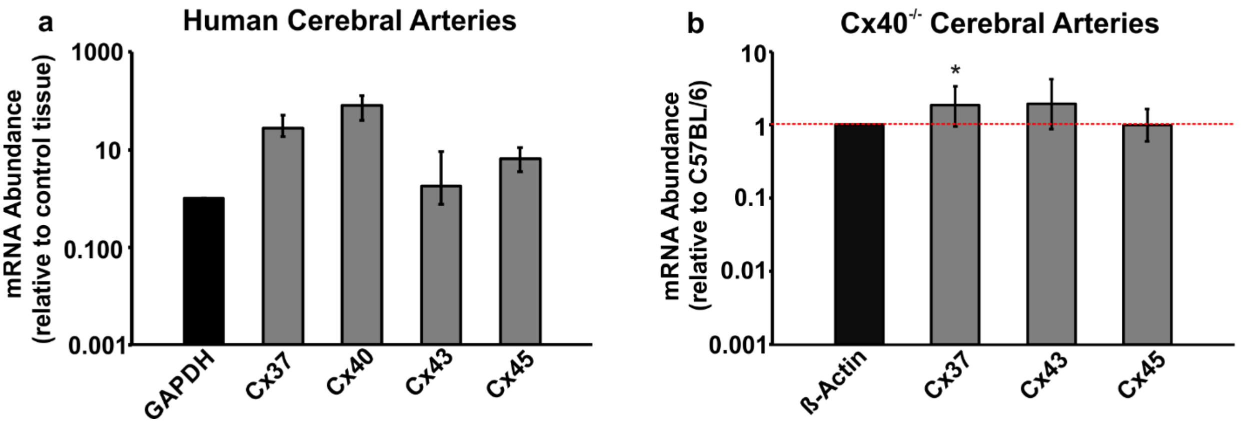


**Figure 6. Vascular connexin (Cx) abundance in human and Cx40-/- mice cerebral arteries. (a)** qPCR analysis revealed the presence of Cx37, Cx40, Cx43 and Cx45 in isolated human cerebral arteries where Cx40 mRNA was observed to be the most abundant. Samples were standardized to reference gene (GAPDH) and were normalized to control tissue (human whole heart RNA). **(b)** A statistically significant increase in Cx37 mRNA was noticed in Cx40-/- mice middle cerebral arteries (MCA), compared to wild type controls (C57BL/6, Cx40+/+). Regulation of other connexins were unaffected by the deletion of Cx40 protein. Samples were standardized to reference gene (ß-Actin) and were compared to MCA from C57BL/6 mice. Red dotted line indicates expression in C57BL/6 cerebral arteries. * denotes significant difference.

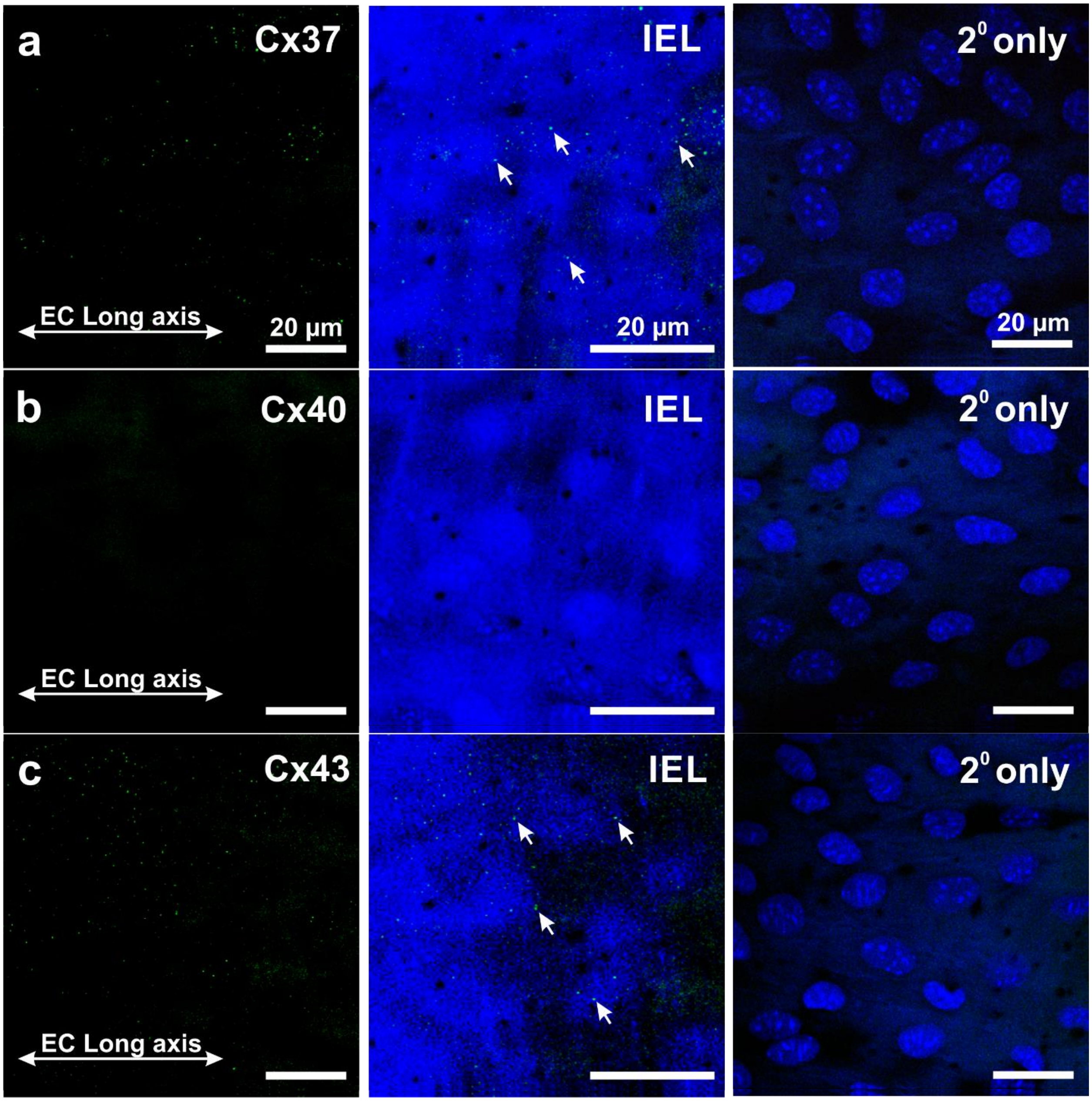


**Figure 7: Endothelial connexin (Cx) expression is diminished in cerebral arteries from Cx40$^{-/-}$ mice. (Left)** Connexin isoforms Cx37 **(a)** and Cx43 **(c)** (green), were identified by immunohistochemistry in the endothelial layer of middle cerebral arteries from Cx40$^{-/-}$ mice. Lack of Cx40 labeling was confirmed in the middle cerebral arteries (**b**). **(Middle)** IEL was delineated by 488 nm autofluorescence and arrowheads denote sites on punctate Cx labeling in the endothelial cell layer. (**Right)** Absence of endothelial Cx labeling in opened arteries incubated without primary antibody; nuclei were labelled with DAPI. Scale bar = 20 µm.

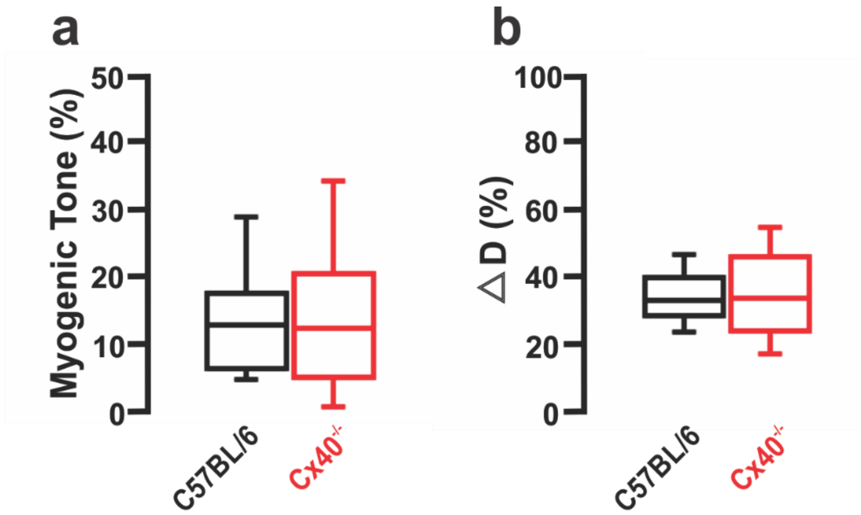


**Figure 8. Cerebral vessel reactivity in C57BL/6 and Cx40$^{-/-}$ mice.** Middle cerebral arteries from mice were isolated and pressurized to 50 mmHg. **(a)** Myogenic tone, expressed as a percent of maximal diameter, was similar in cerebral arteries from C57BL/6 and Cx40$^{-/-}$ mice. Maximal diameter was determined by superfusing cerebral arteries in $Ca^{2+}$ free PSS containing 2 mM EGTA. **b)** KCl (60 mM) was superfused in $Ca^{2+}$ PSS and the constriction responses were monitored. Vasomotor measurements expressed as a percent of resting diameter shows no difference between C57BL/6 and Cx40$^{-/-}$ mice.

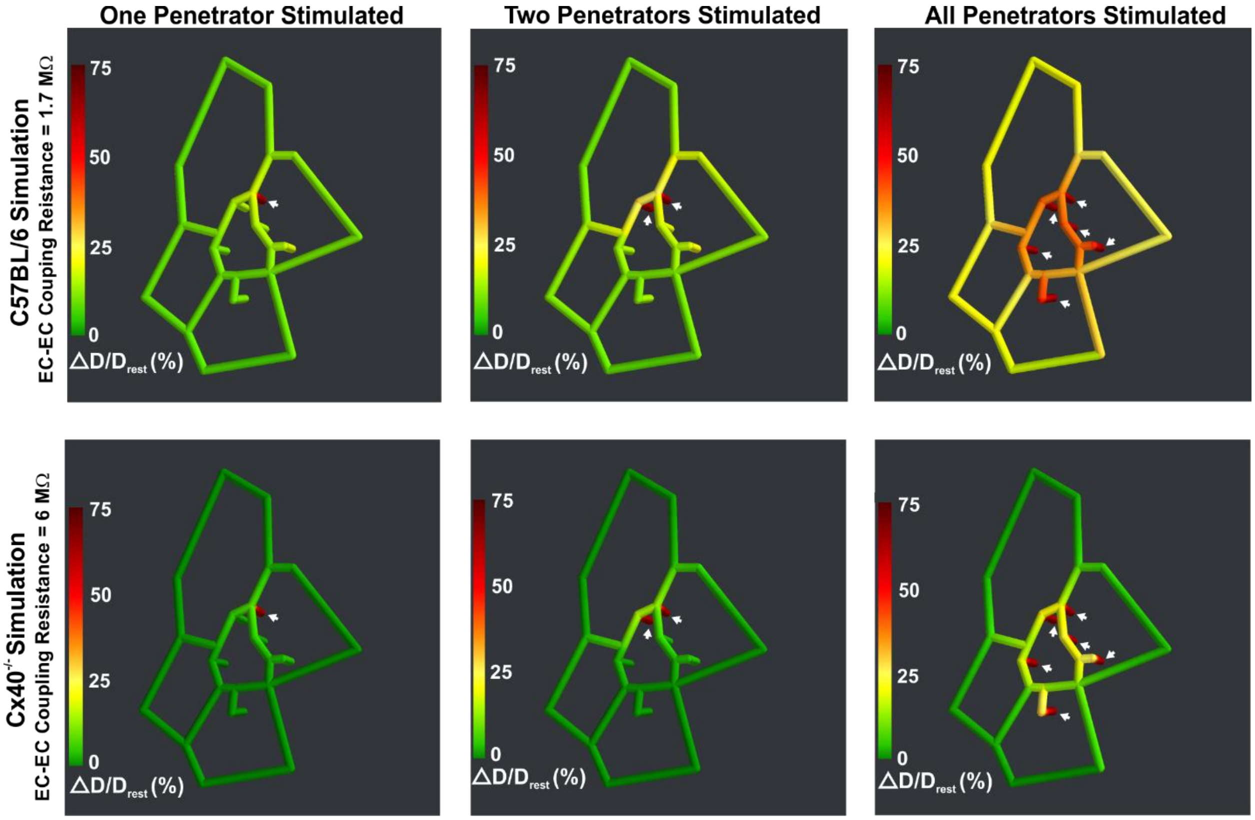


**Figure 9.** ***In silico*** **modeling: The ascension of vasodilation from penetrating arterioles to surface vessels.** Using observations from Shih et al., 2009 (1), a network model of surface arteries and penetrating arterioles was built to study the spread of electrical/vasomotor responses. The *in silico* network consisted of an interconnected network of surface arteries and six adjoining penetrating arterioles. Simulations entailed voltage clamping (15 mV negative to resting $V_m$, 250 ms) one distal segment in one, two or all penetrating arterioles and resolving the vasomotor response throughout the network. The ascension of vasomotor responses was color mapped along the network; endothelial-to-endothelial coupling resistance was set to 1.7 (C57BL/6 control) or 6 (Cx40$^{-}$/$^{-}$) MΩ (calculated in Figure 2). Vasomotor responses were expressed relative to resting diameter.

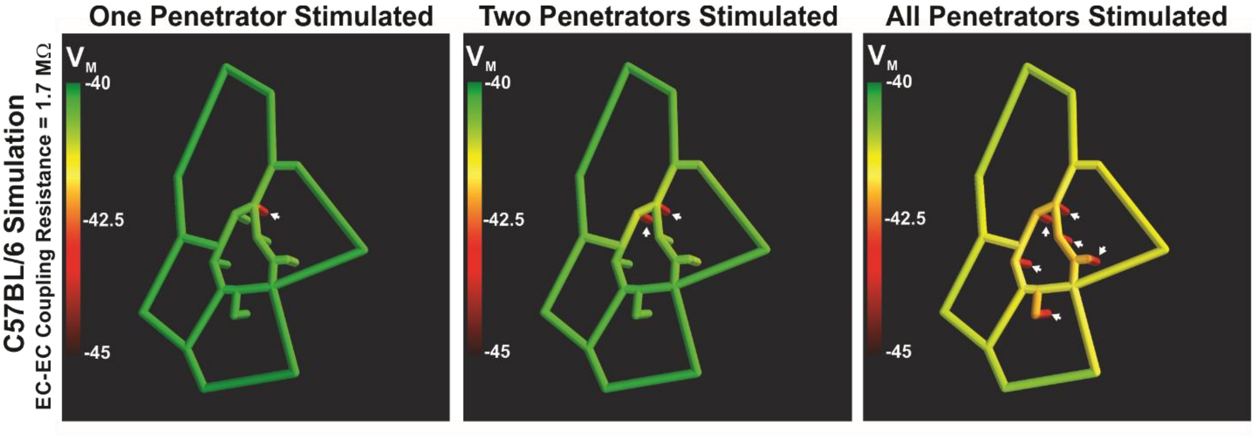


**Figure 10: Ascension of hyperpolarization from penetrating arterioles to surface vessels is preserved at a lower simulation threshold.** Using observations from Shih et al., 2009 (1), a network model of surface arteries and penetrating arterioles was built to study the spread of electrical/vasomotor responses. The *in silico* model consisted of an interconnected network of surface arteries and six adjoining penetrating arterioles. In comparison to Fig. 4, the initial hyperpolarizing response was moderated to 5 mV negative to resting $V_m$ (250 ms). The ascension of electrical responses was color mapped along the network after simulating one, two or all of the penetrators.

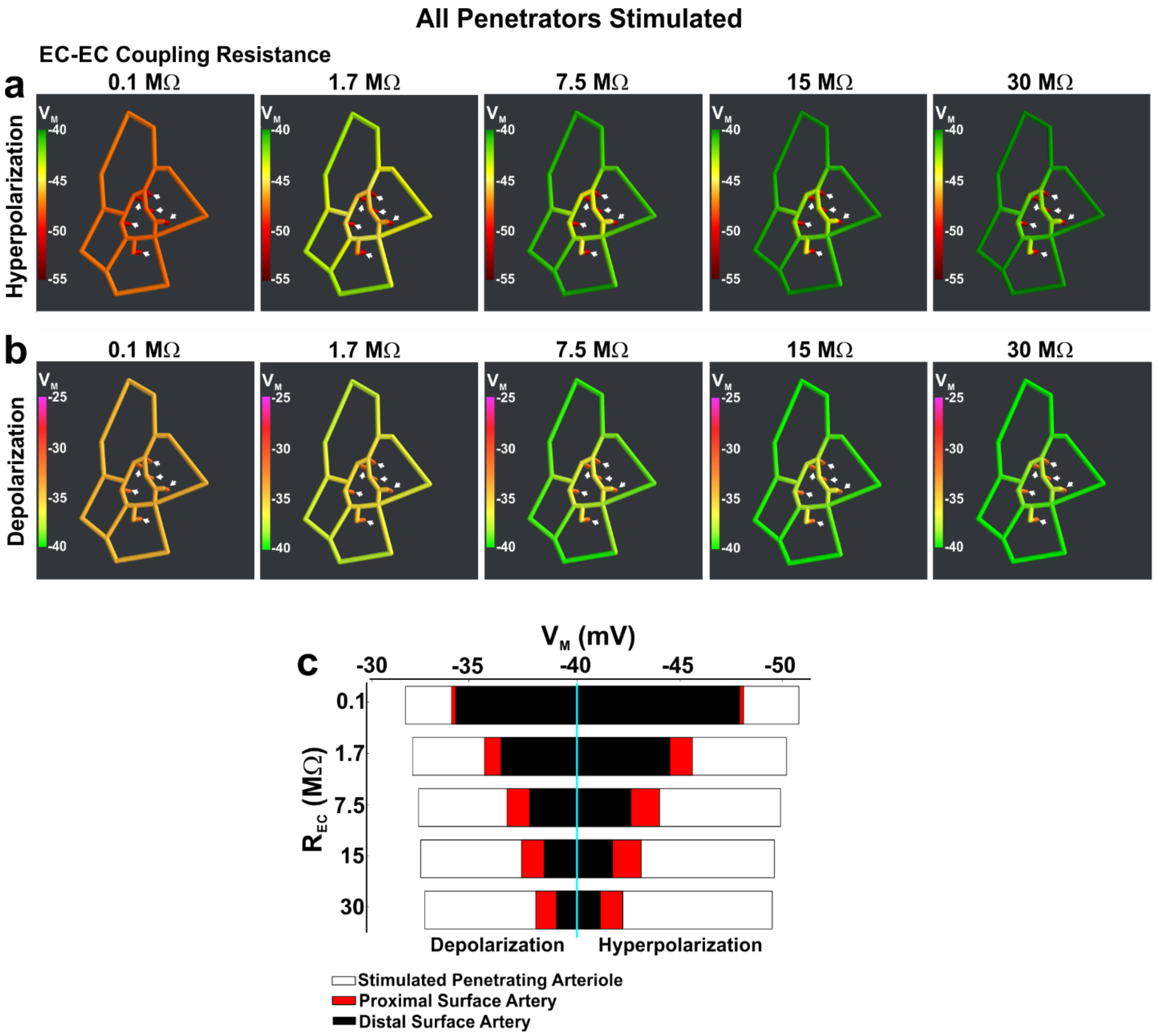


**Figure 11. *In silico* modeling: The impact of endothelial-to-endothelial coupling resistance on the ascension of electrical responses from penetrating arterioles to surface vessels.** Using observations from Shih et al., 2009 (1), a network model of surface arteries and penetrating arterioles was built to study the spread of electrical responses. The *in silico* model consisted of an interconnected network of surface arteries and six adjoining penetrating arterioles. Simulations entailed voltage clamping (15 mV positive or negative to resting $V_m$, 250 ms) one distal segment in all penetrating arterioles and resolving the electrical response throughout the network. **(a, b)** The ascension of electrical responses was color mapped along the network after setting the endothelial-to-endothelial coupling resistance at 0.1, 1.7, 7.5, 15 or 30 MΩ. **c)** Summary data was plotted at 3 sites in the network as defined in Figure 3, which included the penetrating arteriole (200 μm upstream from the stimulation site) and two surface vessel sites.

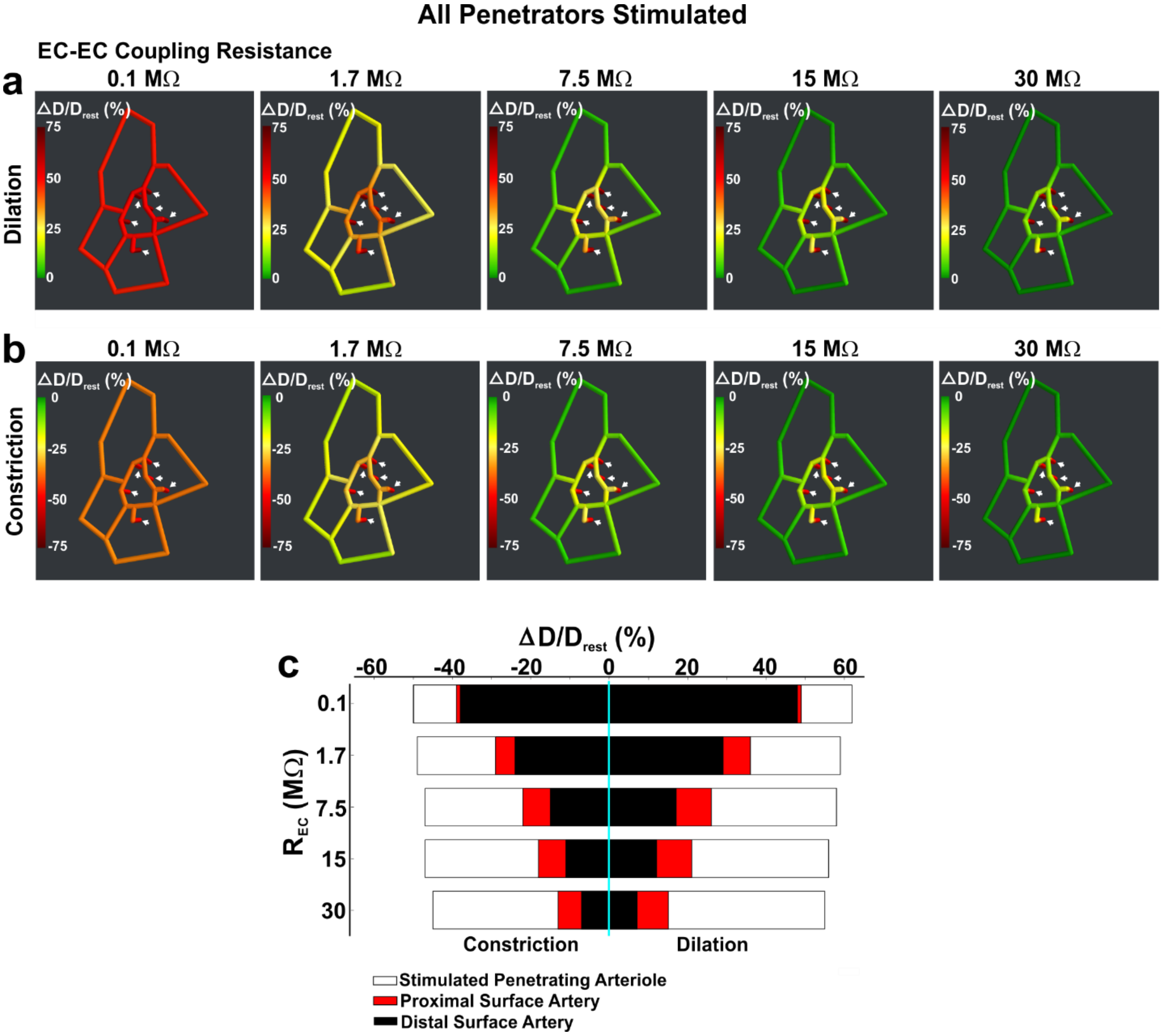


**Figure 12.** ***In silico*** **modeling: The impact of endothelial-to-endothelial coupling resistance on the ascension of vasomotor responses from penetrating arterioles to surface vessels.** Using observations from Shih et al., 2009 (5), a network model of surface arteries and penetrating arterioles was built to study the spread of vasomotor responses. The *in silico* network consisted of an interconnected network of surface arteries and six adjoining penetrating arterioles. Simulations entailed voltage clamping (15 mV positive or negative to resting $V_m$, 250 ms) one distal segment in all penetrating arterioles and resolving the vasomotor response throughout the network. **(a, b)** The ascension of vasomotor responses was color mapped along the network after setting the endothelial-to-endothelial coupling resistance at 0.1, 1.7, 7.5, 15 or 30 MΩ. **c)** Summary data was plotted at 3 sites in the network as defined in Figure 3, which included the penetrating arteriole (200 µm upstream from the stimulation site) and two surface vessel sites. Vasomotor responses were expressed relative to resting diameter.

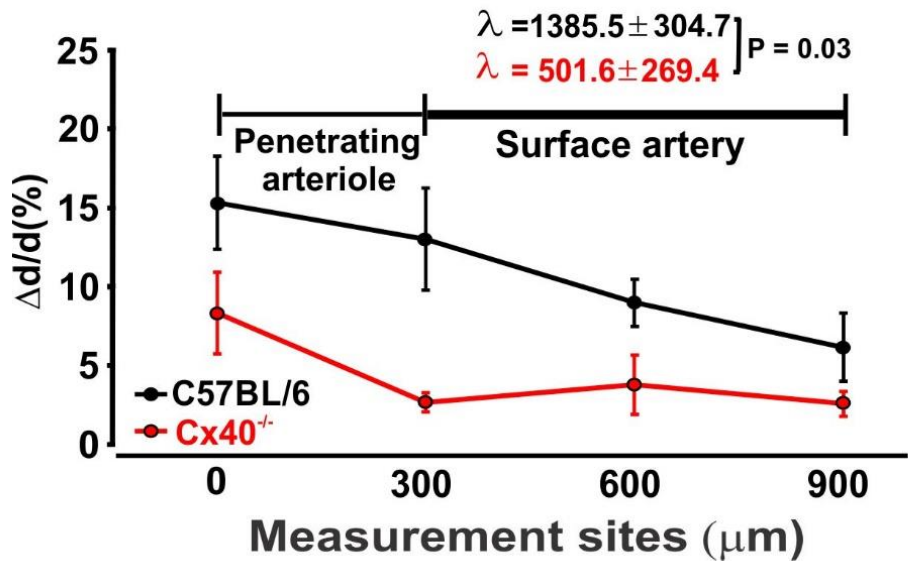


**Figure 13: Compromised vascular signaling impairs conduction of neurovascular coupling responses.** Awake mice were positioned on an exercise ball and the FITC Dextran labelled microvasculature was imaged using two-photon microscopy. Starting near a point of barrel cortex activation vasomotor responses to whisker stimulation were monitored at 4 sites upstream sites (spaced 300 µm apart). To assess conduction decay, individual data sets were fitted to an exponential function f(x) = k*exp(-x/lambda), using the nonlinear least-squares Marquardt-Levenberg algorithm. A single tailed- t-test then compared the summative decay constant from C57BL/6 and $Cx40^{-/-}$ mice. $P < 0.05$ denotes significant difference.

**SI Appendix (Tables)**

**Table 1: Number of animals/vessels per experiment.**

| Results in | **Experiment** | **N number.** |
|---|---|---|
| Figure 1 | Electron Microscopy (C57BL/6)<br>Immunohistochemistry (C57BL/6) | (3)<br>(5) |
| Figure 2 | Human cerebral artery<br>Conduction (Endothelium intact)<br>(C57BL/6)<br><br>($Cx40^{-/-}$)<br><br>Endothelial denuded Conduction<br>(C57BL/6)<br><br>($Cx40^{-/-}$)<br><br>Membrane potential<br>(C57BL/6)<br>($Cx40^{-/-}$) | (450µm, 3; 900µm, 3; 1350µm, 3; 1800µm, 3)<br><br>(0µm, 7; 450µm, 10; 900µm, 10; 1350µm, 8; 1800µm, 8)<br>(0µm, 7; 450µm, 10; 900µm, 10; 1350µm, 10; 1800µm, 10)<br><br>(0µm, 7; 450µm, 7; 900µm, 7; 1350µm, 7; 1800µm, 7)<br>(0µm, 7; 450µm, 7; 900µm, 7; 1350µm, 7; 1800µm, 7)<br><br>(6)<br>(5) |
| Figure 3 | Conducted Dilation<br>Surface Artery<br>Penetrating Arteriole<br>Immunohistochemistry | <br>(Low $K^+$, 4; ACh, 3)<br>(Low $K^+$, 4)<br>(4) |
| Figure 5 | Neurovascular Coupling<br>(C57BL/6)<br><br>($Cx40^{-/-}$) | <br>(0µm, 21; 300µm, 21; 600µm, 19; 900µm, 19: 3 animals)<br>(0µm, 14; 300µm, 14; 600µm, 21; 900µm,16: 6 animals) |
| Figure 6 | Sham Controls<br>(C57BL/6)<br>($Cx40^{-/-}$)<br>Stroke MRI<br>(C57BL/6)<br>($Cx40^{-/-}$) | <br>(4)<br>(4)<br><br>(9)<br>(10) |
| Figure 7 | Stroke MRI, ASL & T2W<br>(C57BL/6)<br>($Cx40^{-/-}$)<br>Cresyl Violet staining<br>(C57BL/6)<br>($Cx40^{-/-}$) | <br>(9)<br>(9)<br><br>(9)<br>(8) |

**Table 2: Number of animals/vessels per experiment.**

| Results in | Experiment | **N Number.** One vessel per animal unless otherwise stated**.** |
|---|---|---|
| SI Appendix<br>Figure 1 | Immunohistochemistry (Cx40)<br>Human cerebral arteries | (2) |
| SI Appendix<br>Figure 2 | Immunohistochemistry<br>Cx37<br>Cx40<br>Cx43 | <br>(4)<br>(4)<br>(4) |
| SI Appendix<br>Figure 6 | qPCR analysis<br>Human cerebral arteries<br>(C57BL/6)<br>(Cx40$^{-}$/$^{-}$) | <br>(6)<br>(6)<br>(7) |
| SI Appendix<br>Figure 7 | Immunohistochemistry<br>Cx37<br>Cx40<br>Cx43 | <br>(4)<br>(4)<br>(4) |
| SI Appendix<br>Figure 8 | C57BL/6)<br>(Cx40$^{-}$/$^{-}$) | (10)<br>(10) |
| Text | Blood Pressure/Heart Rate<br>(C57BL/6)<br>(Cx40$^{-}$/$^{-}$) | <br>(10)<br>(10) |

**Table 3: qPCR-primers utilized for gene expression analysis.**

| Gene Target | Ascension Number | Primer sequence |
|---|---|---|
| Cx37 (GJA4) | NM_008120 | GGT CGT CCC CTC TAC CT<br>ACC GTT AAC CAG ATC TTG CC |
| Cx40 (GJA5) | NM_001271628 | GTT TCA ACT TCG ACC TCA CTC T<br>GCT CCA GTC ACC CAT CTT G |
| Cx43 (GJA1) | NM_010288 | CCT TTG ACT TCA GCC TCC AA<br>GAC CTT GTC CAG CAG CTT C |
| Cx45 (GJC1) | NM_008122 | GGT AAC AGG AGT TCT GGT GAA<br>TCG AAA GAC AAT CAG CAC AGT |
| (β-Actin) | NM_007393 | GGC TGT ATT CCC CTC CAT CG<br>CCA GTT GGT AAC AAT GCC ATG T |

**Table 4: Primary and secondary antibodies used.**

| **Antibody (Ab)** | **Raised against amino acids (aa)/ antigen (Ag, species / epitope)** | **[Ab]** | **Supplier/ Cat #** | **Host** |
|---|---|---|---|---|
| Cx37 | Mouse C-terminus synthetic peptide | 1:200 | Thermofisher, 40-4200 | Rabbit |
| Cx40 | Rat C-terminus, aa 328-340, intracellular | 1:200 | Alomone, ACC205 | Rabbit |
| Cx43 | Human/Rat C-terminus, aa 363-382 with N-terminally added lysine | 1:2000 | Sigma, C6219 | Rabbit |
| Anti-rabbit (H+L) Secondary Antibody, Alexa Fluor 488 | - | 1:1000 | Thermofisher, A-21206 | Donkey |
| Anti-rabbit IgG (H+L) Secondary Antibody, HRP | - | 1:10000 | Thermofisher 65-6120 | Goat |
